# New Insights into nuclear import and nucleolar localization of yeast RNA exosome subunits

**DOI:** 10.1101/2025.02.25.640171

**Authors:** Valdir Gomes Neto, Leidy Paola P. Cepeda, Bruno R.S. Queiroz, Sylvain Cantaloube, Isabelle Leger-Silvestre, Thomas Mangeat, Benjamin Albert, Olivier Gadal, Carla C. Oliveira

## Abstract

The RNA exosome is a multiprotein complex essential for RNA maturation and degradation. In budding yeast, a nine-subunit protein complex (Exo9) associated with Rrp44 forms Exo10 in the cytoplasm and, in complex with Rrp6, Exo11 in the nucleus. Depending on its subcellular localization, the exosome interacts with different cofactors and RNA substrates. In the cytoplasm, Exo10 associates with the SKI complex via Ski7, while in the nucleus, Exo11 interacts with the TRAMP complex. Within the nucleolus, the exosome participates in ribosomal RNA (rRNA) processing, facilitated by Mtr4-dependent adaptors Utp18 and Nop53. In this manuscript, we have performed a comprehensive study that addresses the targeting mechanism and precise subcellular localization of all members of the Exo11 complex. We observed a high concentration of all Exo11 subunits in the nucleolus and identified the importins Srp1 (α) and Kap95 (β) as responsible for the nuclear import of Exo9 subunits. Notably, Exo9 subunits localization was not significantly disrupted in the simultaneous absence of NLS-containing subunits Rrp6 and Rrp44, suggesting redundant nuclear import pathways for Exo9. Additionally, we show evidence that Ski7 may play a role in the Exo9 retention in the cytoplasm. To explore the exosome sub-nucleolar localization, we compared Rrp43 with nuclear exosome cofactors and show that it is enriched in the same nucleolar region as Mtr4 and Nop53. In conclusion, our findings provide a detailed characterization of Exo11 distribution, highlight the primary nuclear import mechanisms for Exo9, and reveal the specific localization of the exosome within the granular component (GC) of the yeast nucleolus, suggesting a spatial regulation of the RNA processing pathway.

**Highlights:**

– Comprehensive study of the localization of all exosome subunits in yeast.
– Identification of karyopherins involved in exosome nuclear import.
– Ski7 dependent cytoplasmic retention of exosome.
– Yeast sub-nucleolar organization of exosome and its cofactors.

## Introduction

In eukaryotes, transcription of rRNA precursors occurs in the nucleolus, a highly organized nuclear compartment with three distinct sub-nucleolar regions, identified by their morphological appearance in transmission electron microscopy and presence of specific nucleolar proteins^1,2^. Transcription by RNA polymerase I occurs on the border between the fibrillar center (FC) and the dense fibrillar component (DFC), from where the nascent pre-rRNA is cotranscriptionally directed to the DFC, and where the early processing reactions start by association of the pre-rRNA with early binding ribosomal proteins and assembly factors. The intermediate processing reactions take place at the granular component (GC), followed by late and final maturation steps in the nucleoplasm and cytoplasm, respectively^3,4^.

Ribosome biogenesis involves the coordinated transcription, modification, processing, and surveillance of precursor rRNAs, which undergo several exo- and endonucleolytic cleavage reactions during their maturation process^5,6^. In the pre-rRNA maturation pathway, the RNA exosome complex is responsible for the degradation of the spacer sequence 5’-ETS, released co-transcriptionally after cleavage at A_0_, and for the 3′-to-5′ end processing of 7S pre-rRNA to form the mature 5.8S rRNA (Fig. S1)^7–9^. The SSU processome is also formed co-transcriptionally by association of U3 snoRNP and other factors to the pre-rRNA that participate in 40S maturation^10^. Later in pre-rRNA processing, endonucleolytic cleavage at C_2_ site in ITS2 of pre-rRNA 27S separates the 7S (5.8S + 5’ region of ITS2) and 26S (3’ region of ITS2 plus 25S) pre-rRNAs^11^, which undergo exonucleolytic processing by the exosome and Rat1/Rai1, respectively, to generate mature rRNAs 5.8S and 25S^11,12^. Mtr4 and the RNA exosome are essential for the 5′-ETS degradation and for ITS2 processing of the 7S pre-rRNA after the cleavage at C_2_^12^, when the exosome subunit Rrp44 shortens 7S to the intermediate 5.8S+30, which is then handed over to Rrp6, responsible for trimming it to 6S pre-rRNA. This is subsequently further processed in the cytoplasm to produce the mature 5.8S rRNA^13^. Exosome is also involved in quality control steps of rRNA processing, targeting unprocessed 35S rRNA and 23S rRNA for degradation, the latter generated by direct cleavage at A_3_ site^14–16^.

In *Saccharomyces cerevisiae*, the exosome is composed of a nine-subunit core (Exo9) that contains a heterohexameric ring formed by the RNase PH domain-containing subunits Rrp41, Rrp42, Rrp43, Rrp45, Rrp46 and Mtr3, and a heterotrimeric “cap” formed by the RNA binding subunits Rrp4, Rrp40 and Csl4. Although the structure of the exosome core is conserved from archaea to eukaryotes, it has phosphorolytic RNase activity in archaea, but no catalytic activity in eukaryotes^17^. In yeast, Exo9 interacts with Rrp44/Dis3 in the nucleus and cytoplasm to form a 10-subunit complex (Exo10). Rrp44 is an RNase II family member, and has two catalytic sites, one with endoribonucleolytic activity (PIN) and a second with processive 3′-to-5′ exoribonucleolytic activity (RNB)^18–20^. The nuclear exosome contains the additional subunit Rrp6 (forming Exo11), an extra catalytic subunit with a distributive 3′-to-5′ exoribonuclease activity that binds to the trimeric cap and upper portion of the hexameric ring, opposite to the Rrp44 binding site^21^. Rrp6 is the only nonessential exosome subunit, although the deletion of its gene results in a slow growth phenotype, temperature sensitivity, filamentous growth and RNA processing defects^7,22–26^.

Although the structure and function of the exosome have been extensively studied in recent years, detailed information on the mechanisms responsible for the subcellular localization of its different forms is still lacking. We have previously identified the nuclear localization signals (NLS) of Rrp6 and Rrp44 and the importins involved in their nuclear transport^25,27^. In addition, we have recently shown that the exosome complex is highly concentrated in the nucleus, and more specifically in the nucleolus^27^, where pre-rRNA is transcribed and where the early processing reactions take place, results that were later supported by the isolation of the 90S pre-ribosome associated with the exosome^28^. To better characterize the subcellular localization of the exosome, here we analyzed all eleven subunits, and show that, consistent with previous results, the complex is concentrated in the nucleolus, where it can readily process pre-rRNA, degrade the spacer 5’-ETS as soon as it is released, as well as misprocessed intermediates such as 23S. We have previously shown that the catalytically active exosome subunits are independently not responsible for signaling the nuclear localization of the remaining subunits, leaving open the question of the mechanism for transport of the exosome core to the nucleus, which we address here and show the involvement of the karyopherins Srp1 and Kap95 in the nuclear import of the exosome catalytically inactive subunits. In addition, we show the importance of the exosome cofactors for its subcellular localization, highlighting the role of Ski7 in the retention of the exosome in the cytoplasm. We address further the localization of the exosome and show for the first time that it is concentrated in the granular component of the nucleolus. Furthermore, our data confirm the presence of three subnucleolar compartments in yeast, similar to what has been described for metazoans.

## Results

### All eleven subunits of exosome are concentrated in yeast nucleolus

One of the essential functions of the exosome in yeast is the processing of pre-rRNA in the nucleus, which underlines the relevance of uncovering the mechanism of nuclear import that allows this protein complex to enter the nucleus. We have previously shown that the exosome catalytically active subunits, Rrp6 and Rrp44, are transported to the nucleus by the karyopherins Srp1/Kap95 or Sxm1 (in the case of Rrp6)^25,27^, but information on the other nine exosome subunits nuclear transport is still lacking.

Since the catalytically active subunits of the exosome are concentrated in the nucleolus, we decided to analyze the subcellular localization of all subunits fused to GFP by super resolution spinning-disk laser scanning confocal microscope, expressed at endogenous levels. Endogenously fused to mCherry Rpa190 (Rpa190-mCherry), the largest RNA polymerase I subunit, was expressed in the same strains to be used as a nucleolar marker. It should be noted that all strains bearing tagged version of Exo11 grow like wild-type yeast (Fig. S2). The results show that despite being involved in processing, degradation and quality control of all classes of RNA in the nucleus and in the cytoplasm, the exosome complex is concentrated in the nucleolus, underscoring the importance of the exosome for pre-rRNA processing (Fig. 1). Quantification of the localization results establishes the concentration of all exosome subunits in the nucleolus (Fig. 2A), suggesting that the complex as a whole is transported to the nucleus and directed to the nucleolus. Concentration of the exosome subunits in the nucleolus was inferred by the calculation of the GFP-fused protein signal in each compartment, corrected by the background and divided by the area, which gives us a clear view of the nucleolus being the compartment with the highest concentration of the exosome. The quantification of total GFP signal shows high similarity among the eleven subunits, despite some small variation in the GFP signal of the proteins being detected (Fig. 2B).

**Figure 1.**
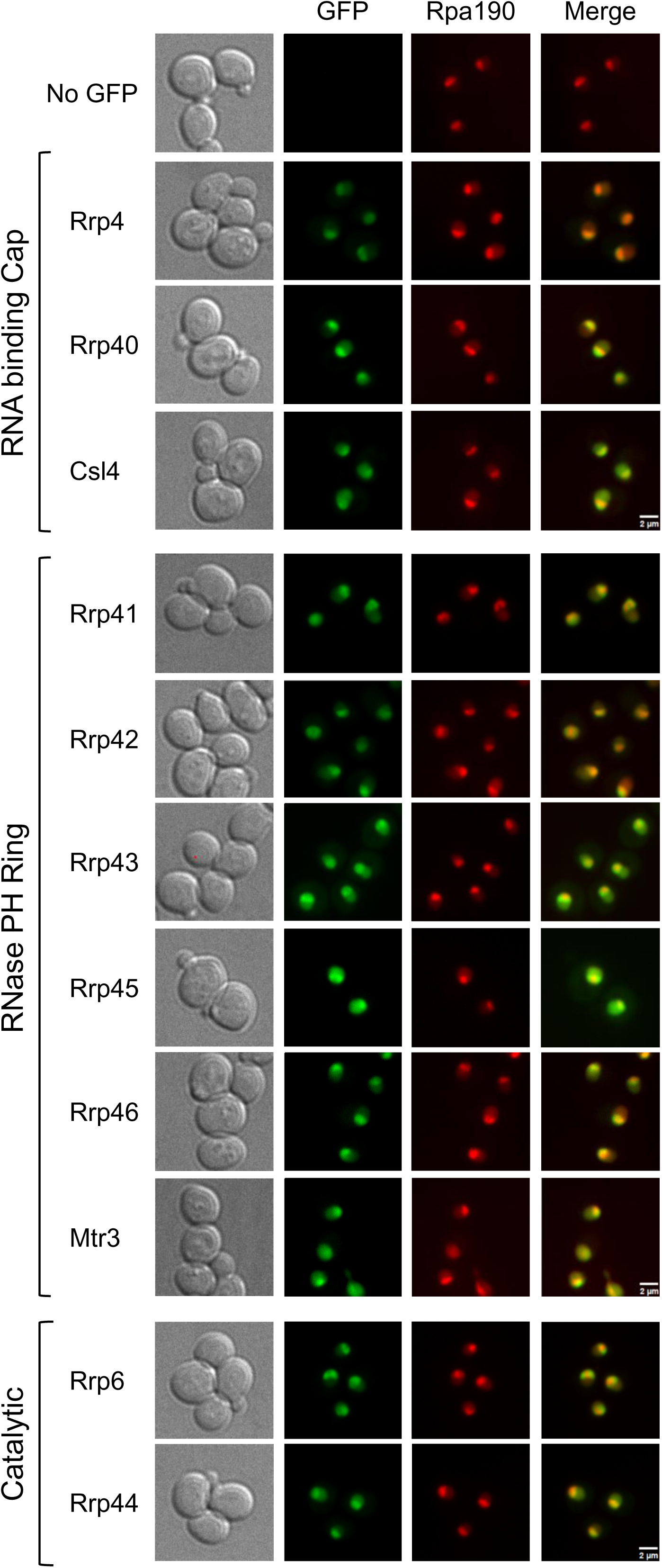
The exosome is mainly nucleolar. Analysis by high-resolution spinning-disk laser scanning confocal microscopy of the subcellular localization of the subunits of the exosome core, fused to GFP and expressed at endogenous levels. Cap, three subunits containing RNA binding domains that form the exosome cap; PH Ring, six subunits with inactive RNase PH domains that form the heterohexameric ring; Catalytic, two catalytically active exosome subunits, which bind the core at opposite sides. Rpa190, fused to mCherry, was used as nucleolar marker. No GFP, control strain expressing only Rpa190-mCherry fusion.

**Figure 2.**
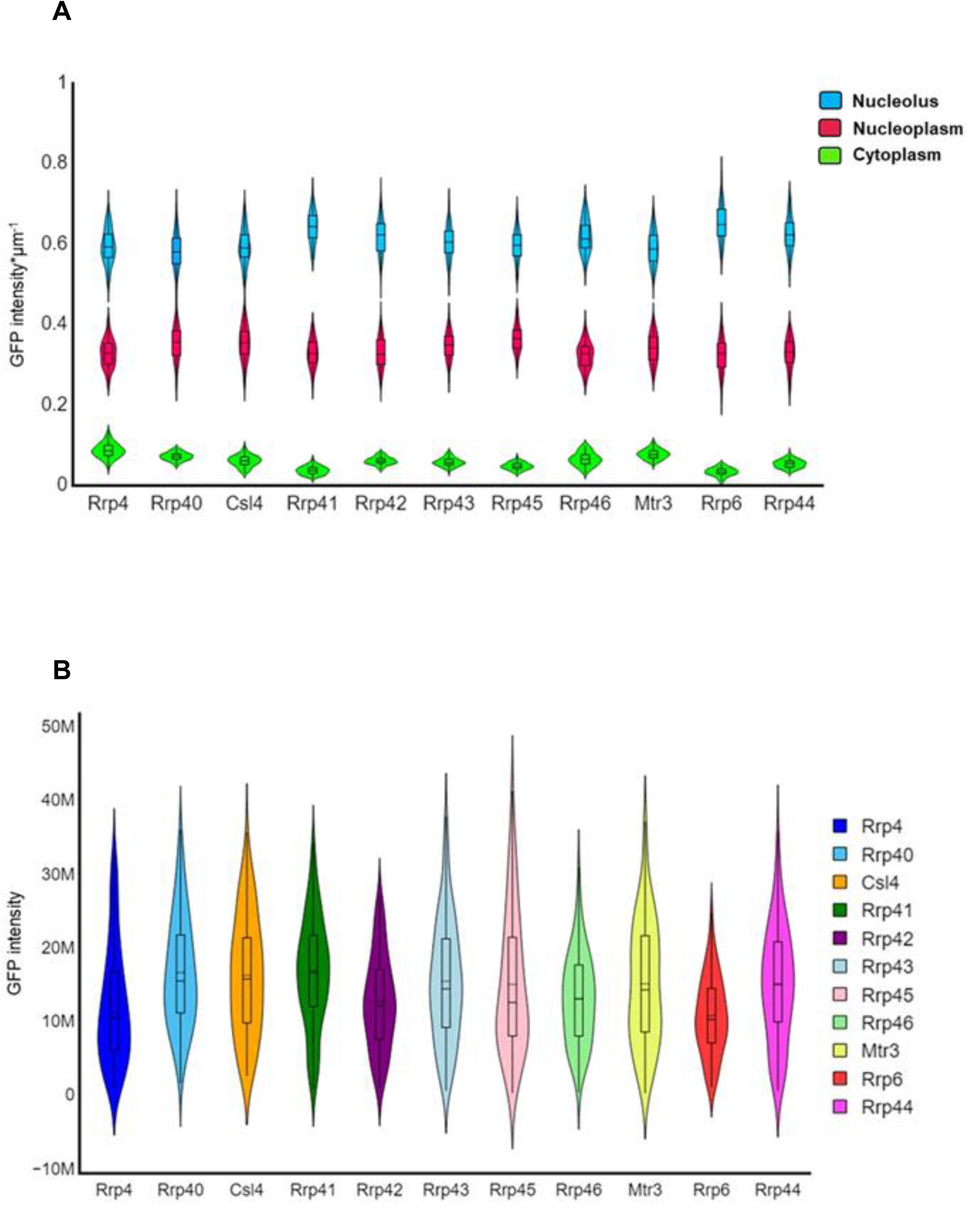
Quantification of the protein signals from the high-resolution spinning-disk laser scanning confocal microscope images in the different subcellular compartments. (**A**) Concentration of GFP-fused RNA exosome subunits in nucleolus, nucleoplasm and cytoplasm. Concentration in each compartment was considered (Intensity of compartment - background)/area. (**B**) Mean intensity of RNA exosome subunits from Z-projection. Total Intensity was considered (Intensity of GFP signal of the entire cell) - (background for an equivalent area).

The yeast nucleolus occupies a small volume of approximately 1μm³, corresponding to 1/50th of the total cell volume^29^. Confirming the exosome concentration in the nucleolus, approximately 20% of the total GFP signals of all subunits were detected within this compartment (Fig. S3A). About 40% of the total GFP signals were in the nucleoplasm (Fig. S3B). While the GFP signal concentration in the cytoplasm was generally lower across all exosome subunits, its relative distribution varied among specific subunits (Fig. 2A). On average, the CAP subunits (Rrp4, Rrp40, and Csl4), the RNase PH Ring subunits (Rrp42, Rrp43, Rrp45, Rrp46, and Mtr3), and the catalytic subunit Rrp44 exhibited approximately 40% of their GFP signals in the cytoplasm. Notably, the core subunit Rrp41 showed a lower signal in the cytoplasm, similar to Rrp6, which is exclusively nuclear (Fig. S3C).

To ascertain that these GFP signals and compartment areas quantifications were performed in similar cells, the cells areas were determined, as well as the areas of the subcompartments, which show that they were very uniform among the strains expressing the exosome subunits (Fig. S4), confirming our conclusions of exosome concentration in the nucleolus.

### Effect of the depletion of the karyopherins on nuclear import of the exosome core subunits

In order to better characterize the nuclear import pathway of all the exosome core subunits and to identify the karyopherins involved in this process, the exosome core subunits were endogenously GFP-tagged in strains in which the genes of either the karyopherin *SRP1* or *KAP95* were placed under control of the *GAL1* promoter, allowing their depletion in glucose-containing medium (Fig. S5). Depletion of Srp1 or Kap95 strongly affects GFP-core exosome localization, leading to partial mislocalization of these proteins to the cytoplasm, albeit still remaining concentrated in the nucleus (Figs. 3 and 4). These results indicate that karyopherins Srp1 and Kap95 are involved in nuclear import of the exosome. Interestingly, while the localization of all eleven subunits is similarly affected by the depletion of Srp1, with the proteins being visualized in higher concentration in the cytoplasm in glucose medium (Fig. 3), depletion of Kap95 did not affect the localization of Csl4 to the same extent (Fig. 4A), which brings a new layer of complexity to the transport of the exosome.

**Figure 3.**
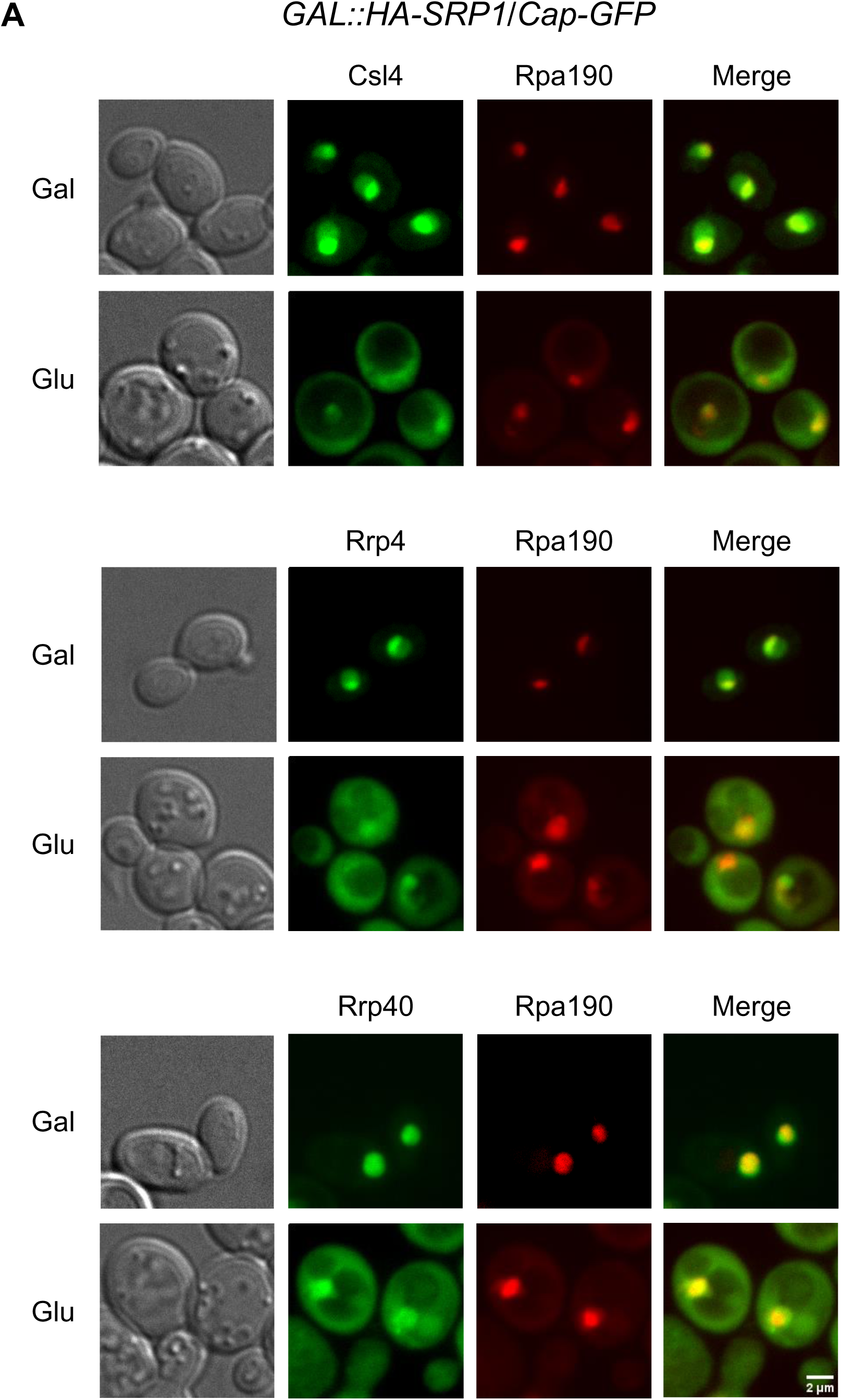
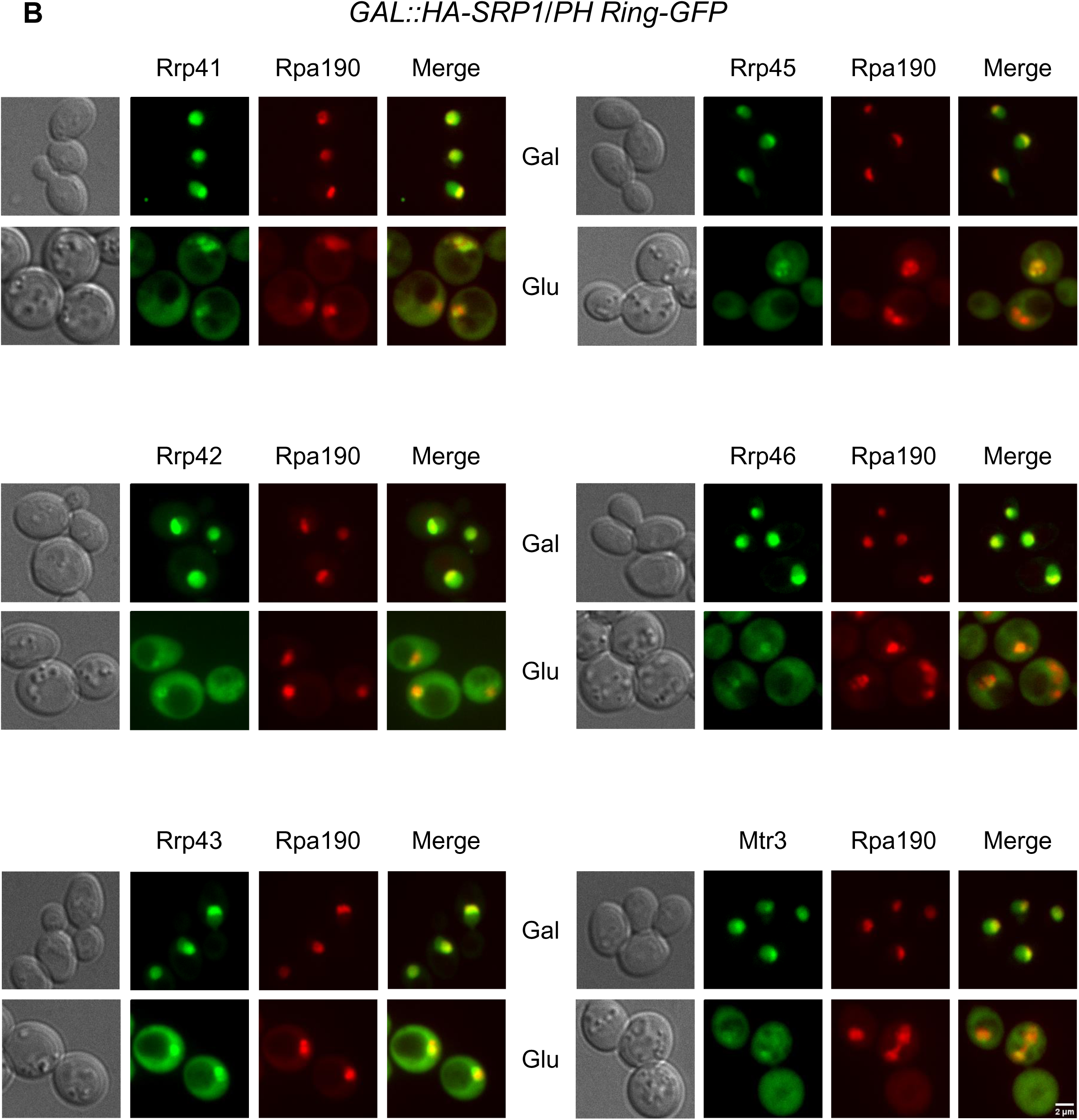
Srp1 affects the nuclear import of the core exosome subunits. Localization of the subunits fused to GFP expressed in *GAL1::HA-SRP1* was analyzed by Z projection in spinning disk confocal microscope, showing that depletion of Srp1 (glucose) affects the exosome core nuclear import. RPA190-mCherry was used as a nucleolar marker. (**A**) Analysis of the exosome cap subunits fused to GFP, Csl4, Rrp4 and Rrp40. (**B**) Analysis of the RNase PH ring subunits, Rrp41, Rrp42, Rrp43, Rrp45, Rrp46 and Mtr3.

**Figure 4.**
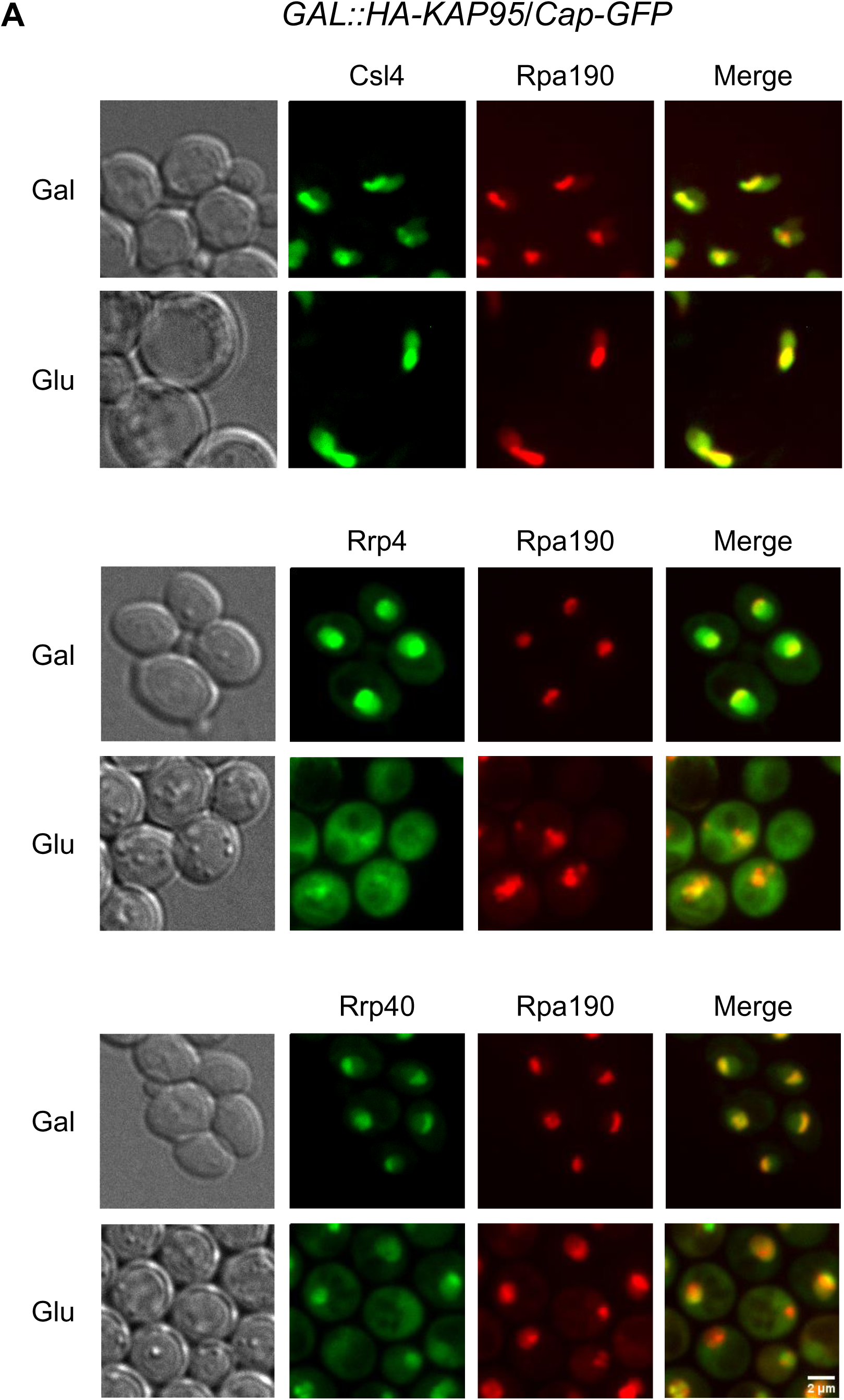
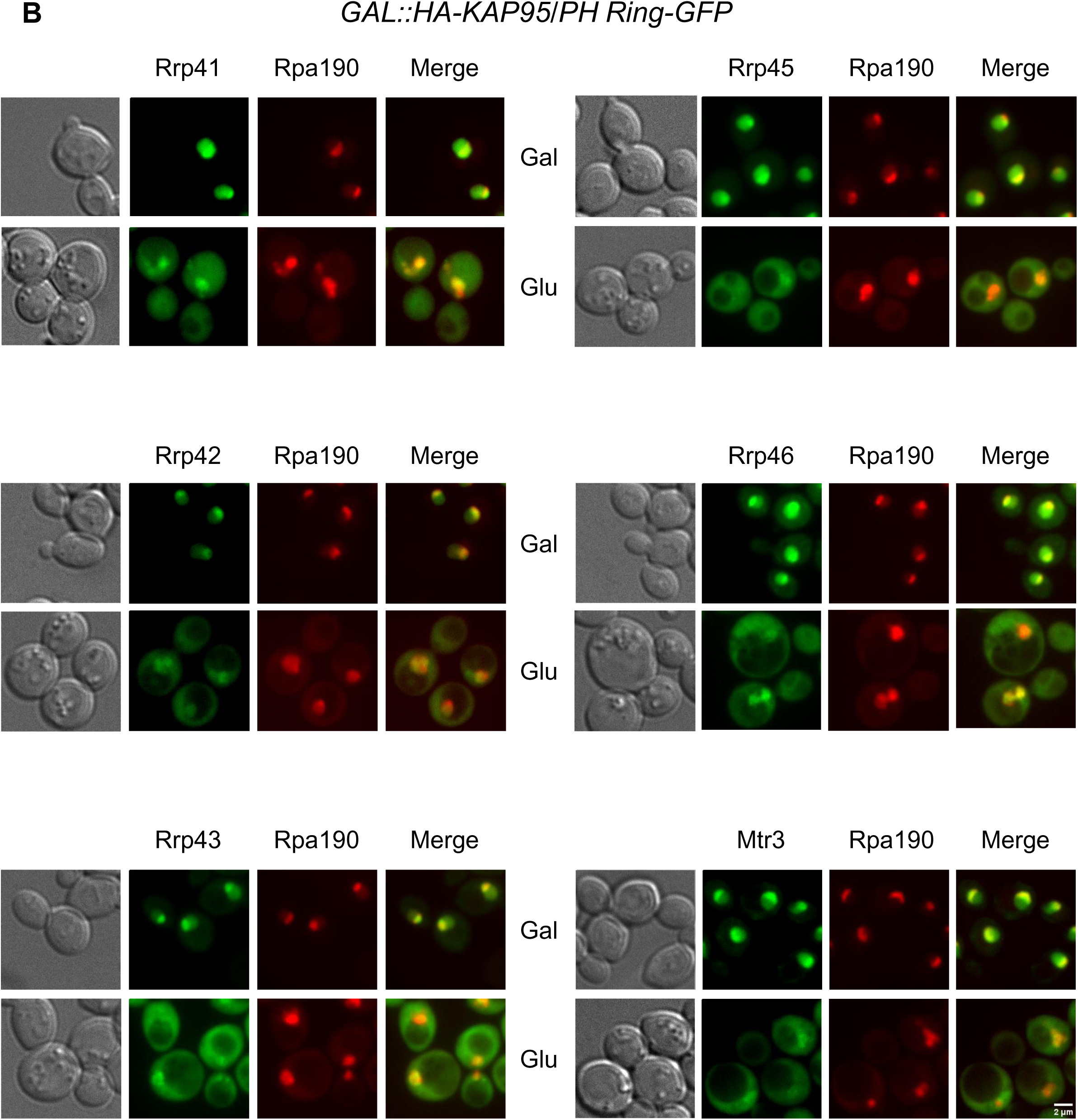
Kap95 affects the nuclear import of the core exosome subunits. Localization of the subunits fused to GFP expressed in *GAL1::HA-KAP95* was analyzed by Z projection in spinning disk confocal microscope, showing that depletion of Kap95 (glucose) affects the exosome core nuclear import. RPA190-mCherry was used as a nucleolar marker. (**A**) Analysis of the exosome cap subunits fused to GFP, Csl4, Rrp4 and Rrp40. (**B**) Analysis of the RNase PH ring subunits, Rrp41, Rrp42, Rrp43, Rrp45, Rrp46 and Mtr3.

To investigate the possible involvement of other β-importins in the nuclear transport of Csl4, the localization of Csl4-GFP was also tested in the deletion strains *Δsxm1*, *Δkap123* and *Δmsn5* (Fig. S6). Surprisingly, Csl4 remained concentrated in the nucleolus in these mutant strains, colocalizing with Rpa190 (Fig. S6A). The localization of Rrp43-GFP was also analyzed in these mutant strains, displaying a nucleolar concentration similar to Csl4 (Fig. S6B). As shown above, Srp1 and Kap95 are involved in the transport of Rrp43 (Figs. 3 and 4), so that no, or little effect of the depletion of other karyopherins was expected in this case. Although the exact β-importin, acting alone or through redundant NLSs recruiting more than one β-importin, involved in the transport of Csl4 remains to be identified, this result raises the question of whether the exosome is transported to the nucleus as a whole core complex or in the form of sub-complexes.

### Depletion of Rrp43 affects nuclear concentration of other exosome core subunits

Contrary to Rrp6 and Rrp44, the Exo9 subunits do not have canonical nuclear localization signals (NLS), with the exception of Rrp43, which contains a putative NLS embedded in the inactive RNase PH domain between the residues 203 and 213 (LKMKRKWSYVL). However, based on the structure of the yeast exosome^21^, this portion of Rrp43 would be only partially exposed, hindering its recognition by importins. Surprisingly, therefore, depletion of Rrp43 affects the nucleolar localization of Exo9 subunits. RNase PH ring subunits Rrp45, Rrp46 and Mtr3, fused to GFP, were expressed at endogenous levels in the strain *GAL1::RRP43/RPA190-mCherry*, incubated in galactose- or glucose-containing medium. The results show that depletion of Rrp43 for six hours in glucose affects the localization of these three exosome subunits, which become visible in the cytoplasm, despite remaining concentrated in the nucleolus (Fig. 5; Fig. S7). Intriguingly, longer periods of time in glucose instead of increasing the GFP signal in the cytoplasm, lead to lower signals in the whole cells, indicating that these exosome subunits might be less stable in the absence of Rrp43, probably due the fact that the core complex is not being formed. This hypothesis was confirmed by the analysis of the levels of these three RNase PH ring components upon depletion of Rrp43. Western blots show decreasing levels of Rrp45, Rrp46 and Mtr3 over time in glucose medium (Fig. S7C).

**Figure 5.**
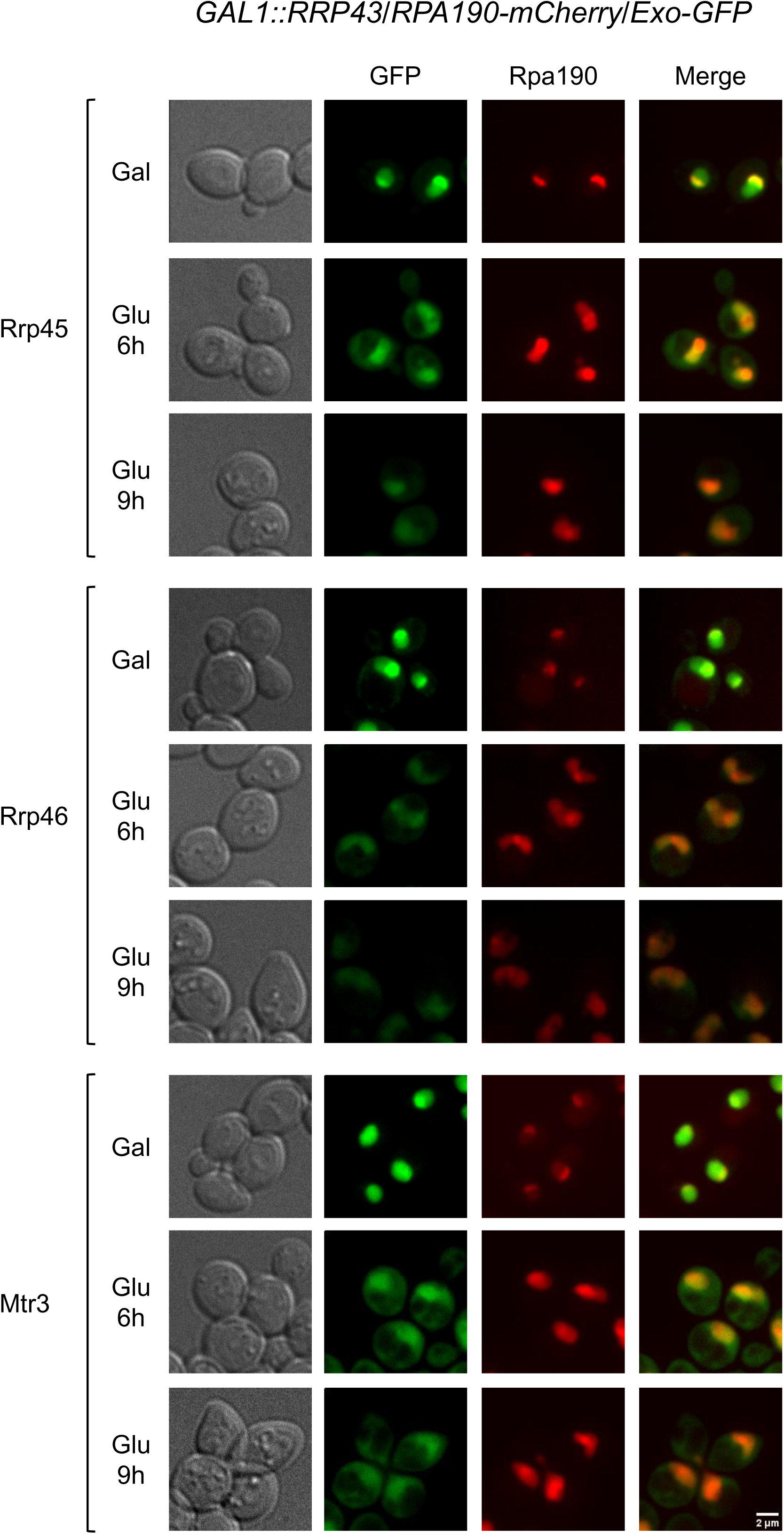
Rrp43 affects the localization of the RNase PH ring exosome subunits. Analysis by high resolution confocal microscopy of the subcellular localization of the RNase PH ring subunits of the exosome core, Rrp45, Rrp46 and Mtr3, fused to GFP and expressed at endogenous levels in the strain *GAL1::RRP43/RPA190-mCherry*, grown in galactose or glucose-containing medium for 6h or 9h. Rpa190-mCherry, was used as nucleolar marker.

To further investigate the nuclear import of the exosome core, we tested the effect of the absence of both exosome catalytic NLS-bearing subunits on the localization of the core subunits. The subcellular localization of the endogenously GFP-fused subunits Mtr3, Rrp43 and Rrp40 were analyzed in the strain *Δrrp6/GAL::RRP44/RPA190-mCherry*. Absence of Rrp6 and depletion of Rrp44 by shift to glucose medium had little effect on the concentration of Mtr3 in the nucleolus, although a stronger GFP signal could be visualized in the nucleoplasm and cytoplasm (Fig. 6A). The same pattern was seen for Rrp43 and Rrp40, which continued concentrated in the nucleolus, but showed higher levels in the cytoplasm in glucose medium (Fig. 6B, C), confirming what we have observed for episomal Rrp41-GFP and Rrp43-GFP in *GAL::RRP44* strain^27^. These results, together with those of the depletion of Rrp43 (Fig. 5), suggest that the exosome is transported to the nucleus as a whole complex, since disruption of the core by the depletion of one of its subunits, affects the nucleolar concentration of the complex.

**Figure 6.**
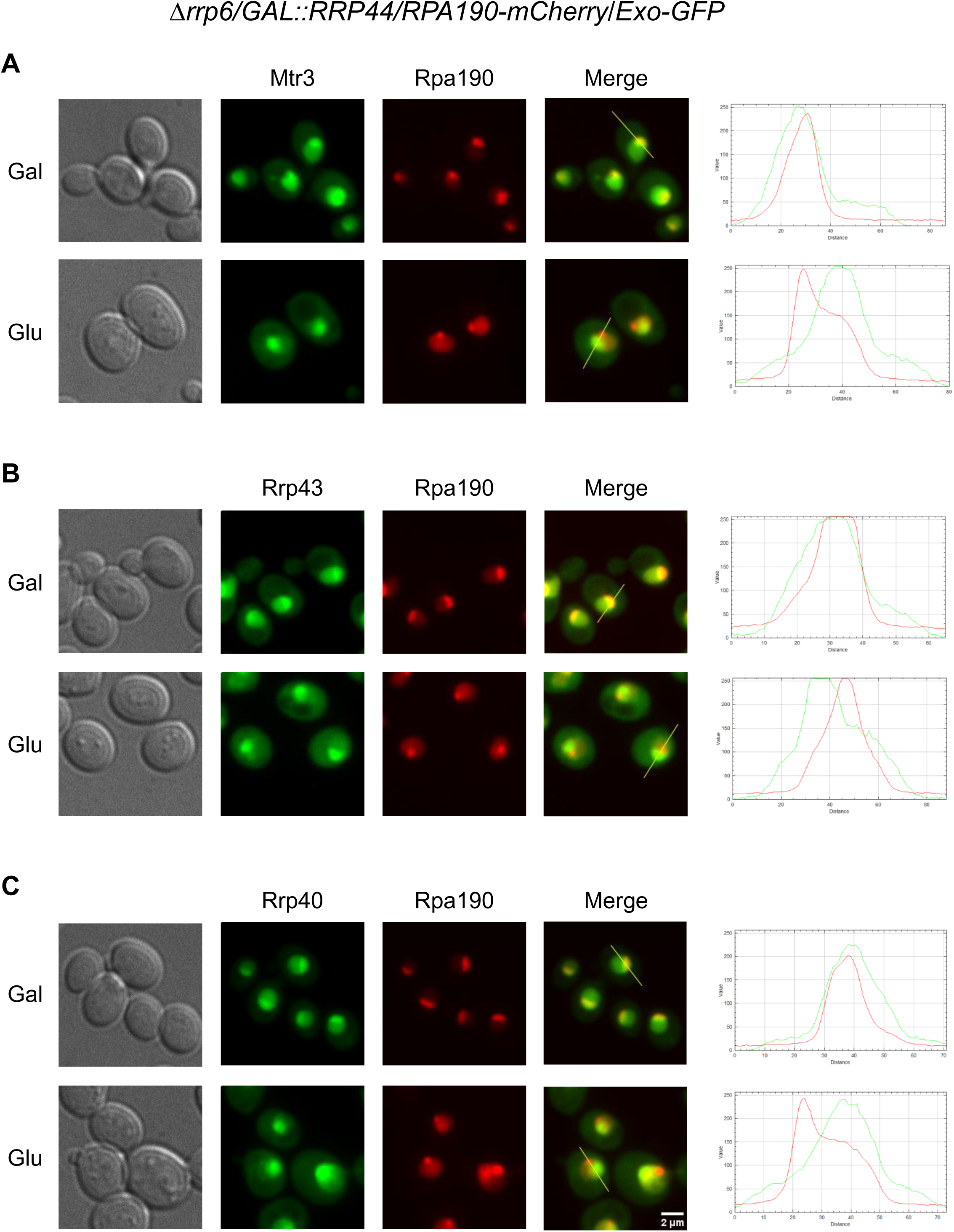
Exosome catalytic subunits affect mildly core subunits localization. Localization of core subunits was analyzed by Z projection in spinning disk confocal microscope in the strain *Δrrp6/GAL::RRP44/RPA190-mCherry* in galactose (presence of Rrp44) or in glucose medium for 6 hours (depletion of Rrp44). (**A**) Mtr3-GFP (**B**) Rrp43-GFP (**C**) Rrp40-GFP. RPA190-mCherry was used as a nucleolar marker. ImageJ plot is shown on the right, where green line represents GFP and red, RPA190-mCherry.

Surprisingly, however, the localization of Rpa190, which was used as a nucleolar control, was affected by the absence of Rrp44 (Fig. 6). With the depletion of Rrp44 in glucose medium, the Rpa190-mCherry signal, despite remaining concentrated in the nucleolus, was also visible in the nucleoplasm. This effect can be seen both by Z projection (Fig. 6) or Z section (Fig. S8) in spinning disk confocal microscope. This result might be due to an effect of the exosome on rRNA transcription. Accordingly, genetic interaction between the exosome and RNA polymerase I subunits has already been reported, in which an RNA polymerase I mutant with transcription elongation defects was shown to be synthetic lethal with Rrp6 deletion and to lead to the accumulation of unprocessed 35S pre-rRNA^30^. We conclude that despite indirect effect probably linked with RNA accumulation in the cell resulting from the absence of both exosome catalytic subunits, core exosome subunits must contain an unidentified nuclear localization signal on top of those identified in Rrp6 and Rrp44.

### Cytoplasmic pool of exosome is controlled by tethering dependent on Ski7

Despite being concentrated in the nucleolus, the exosome is also present in the nucleoplasm and in the cytoplasm. Given that the main function of the yeast exosome is in pre-rRNA processing and quality control, we hypothesized that to be present in the cytoplasm, the exosome would have to be retained in this cell compartment. To address the mechanism of exosome retention in the cytoplasm, we investigated the possible role of the SKI complex, a known cytoplasmic exosome interactor and cofactor^31,32^. Ski7 is a cytoplasmic adaptor required for bridging the exosome and the SKI complex, formed by Ski2/3/8^32^. To recruit the exosome to its cytoplasmic targets, Ski7 binds directly Csl4, Mtr3 and Rrp43^33^. Rrp6, the nuclear exclusive exosome subunit, binds the exosome core mainly at the top of the RNA binding cap, contacting all three subunits, Rrp4, Rrp40 and Csl4, although its C-terminal portion also contacts Rrp43^34^. With this structural information, it is possible to hypothesize that the exosome core interactions with Ski7 or Rrp6 are mutually exclusive, and therefore might play a role in exosome transport, or retention in either subcellular compartment, cytoplasm or nucleus, respectively. To test this hypothesis, the localization of the endogenously GFP-fused exosome subunits was tested in a *Δski7* strain bearing a plasmid coding for Ski7-TAP under control of the strong inducible promoter of *MET15* gene (*Δski7/pMET15::SKI7-TAP/RPA190-mCherry/Exo-GFP*). When incubated in medium containing methionine, Ski7 expression in the cells is repressed. In medium without methionine, expression is strongly induced, leading to Ski7 overexpression (Fig. S9C), so that we can analyze its effect on the exosome localization. The results show that Ski7 overexpression has no effect on the localization of Rrp6 and Rrp44 (Fig. S9A), the catalytically active exosome subunits that we have shown to have NLS and to be transported to the nucleus independently^25,27^. The exosome core subunits, on the other hand, show increased cytoplasmic concentration upon overexpression of Ski7 (Fig. 7), especially Csl4, Rrp43 and Mtr3, which are the subunits that directly interact with Ski7. These surprising results strongly suggest that Ski7 may retain the EXO9 complex in the cytoplasm, not Exo10. As an additional control, we compared the localization of Csl4 in *Δski7* with that in WT cells and show that in both cases, Csl4 is concentrated in the nucleolus (Fig. S9B), as opposed to the higher concentration of Csl4 in the cytoplasm upon overexpression of Ski7 (Fig. 7).

**Figure 7.**
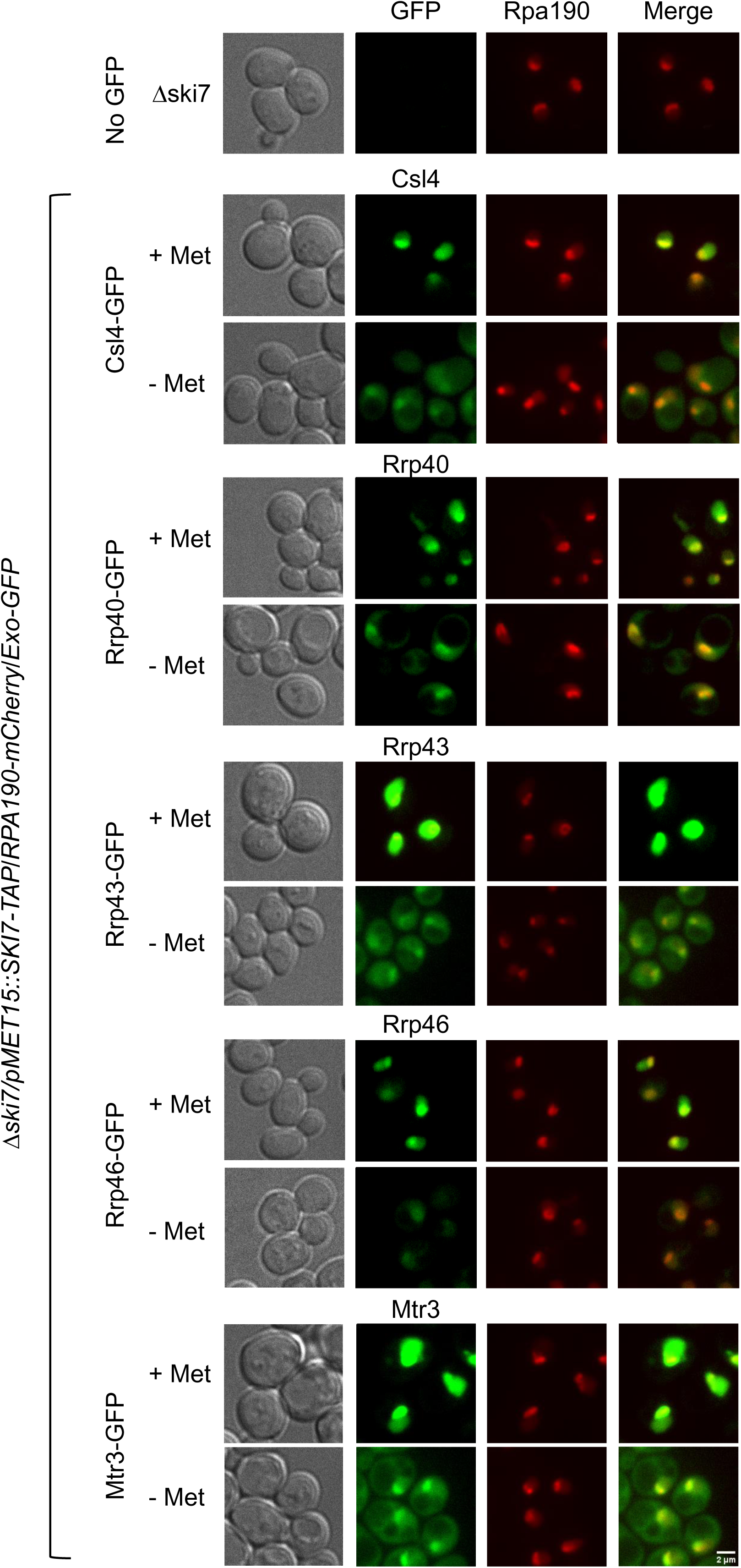
Ski7 affects the localization of the core exosome subunits. Localization of the plasmid-expressed GFP-fused exosome subunits in *Δski7/pMET15::SKI7-TAP/RPA190-mCherry* strain was analyzed by Z projection in spinning disk confocal microscope, showing that overexpression of Ski7 (- Met) affects the exosome core nuclear localization. RPA190-mCherry was used as a nucleolar marker.

### The exosome is concentrated in one subnucleolar area apart from RNA polymerase I

As described above, we have clarified how nuclear import and cytoplasmic retention of the exosome is achieved. However, the localization of the exosome within the nucleolus remains to be better characterized. We have already observed that despite a high concentration in the nucleolus, the exosome subunits do not colocalize exactly with nucleolar RNA polymerase I subunit Rpa190^27^. Interestingly, it has been shown that yeast ribosomal assembly factors localize to distinct nucleolar layers, in which the 40S assembly factors are enriched in an inner and the 60S factors are in an outer layer^35^. Accordingly, in exponentially growing cells, rDNA labelled using the RNA polymerase I activator Net1 fused to mKate is surrounded by snoRNPs^36,37^. To investigate in further detail the exosome nucleolar localization, Rrp43-GFP and the box C/D snoRNP subunit Nop56-GFP were compared to the rDNA marker Net1-mKate in exponentially growing cells, or cells treated with the RNA pol I inhibitor rapamycin^38^ in stationary phase, analyzed on a spinning disk confocal microscope. In agreement with previous data, Net1 showed to be concentrated in a more central area, surrounded by Nop56 in exponential growth, but not upon rapamycin treatment, probably due to the transcription inhibition in the latter (Fig. 8A). Rrp43, on the other hand, shows a different subnucleolar localization relative to Net1 in both growth conditions (Fig. 8B). Net1 is concentrated in a compact area, whereas Rrp43 is spread in an outer layer of the nucleolus relative to Net1 (Fig. 8B). Upon rapamycin treatment, Rrp43 displays a localization similar do Nop56, relative to Net1. However, when directly comparing Rrp43 to Nop56, we show that they do not colocalize in either condition, with Nop56 present in an inner layer than Rrp43 in exponential growth (Fig. 8C), suggesting that the exosome is not concentrated in the same subcompartment as Nop56. Rrp43, however, colocalizes with Rrp1, a 27S pre-rRNA maturation factor^39^, process in which the exosome plays an import role (Fig. 8D)^39^. Although all these proteins localize to the nucleolus, they display different subnucleolar localization. Furthermore, these super-resolution images show the layers in which Net1, Nop56 and Rrp43/Rrp1 are concentrated, confirming the presence of at least three nucleolar subcompartments in yeast, similar to what is seen in metazoan cells.

**Figure 8.**
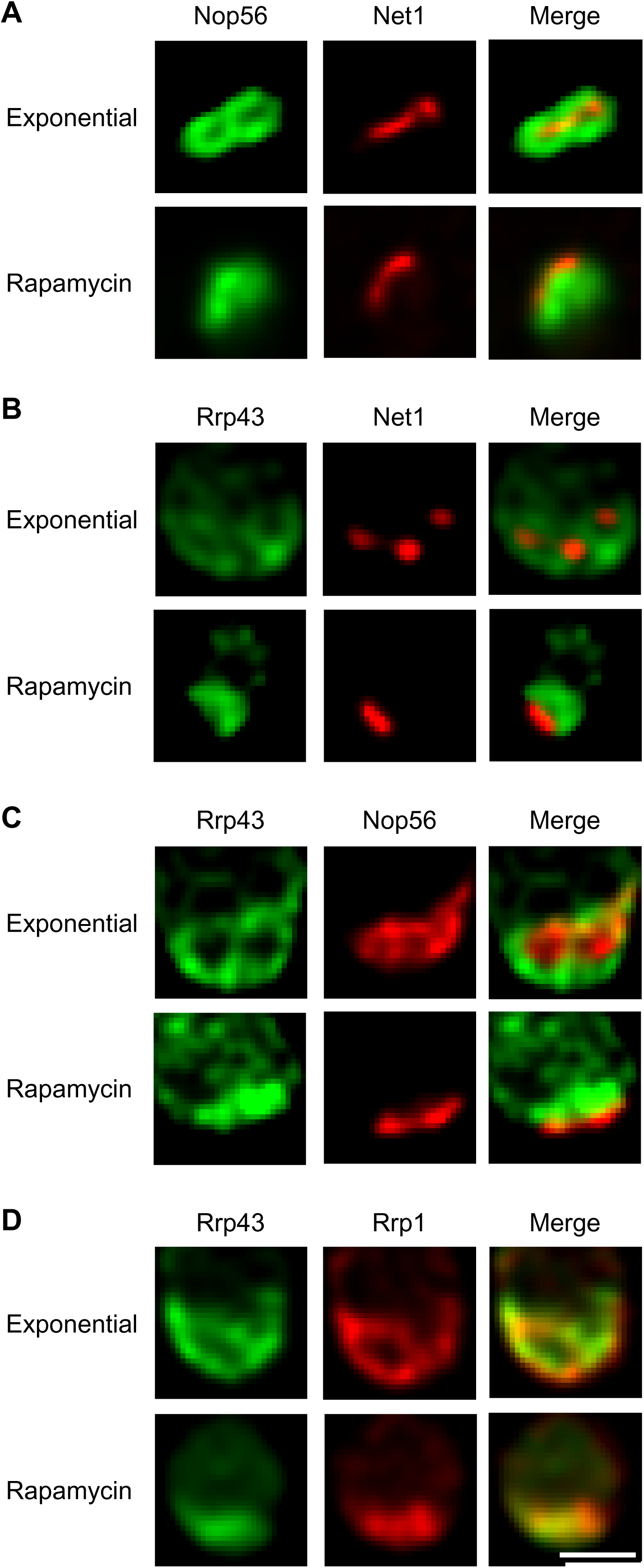
Rrp43 colocalizes with Rrp1, not with Net1 or Nop56 in the nucleolus. Images obtained by Z section of spinning disk confocal microscope of Rrp43-GFP compared to Net1-mKate (rDNA marker) or Nop56-mCherry (DFC marker) in asynchronous cells either on exponential growth or after rapamycin treatment. (**A**) Localization of Nop56-GFP and Net1-mKate. (**B**) Rrp43-GFP and Net1-mKate. (**C**) Rrp43-GFP and Nop56-mCherry. (**D**) Rrp43-GFP and Rrp1-mCherry.

Furthermore, when comparing Rrp43-GFP subnucleolar localization with exosome cofactors, we found that Rrp43 colocalizes with Mtr4 and Nop53, in accordance with the recruitment of the exosome during maturation of pre-60S, when the exosome processes 7S pre-rRNA to give rise to mature 5.8S rRNA (Fig. 9). Interestingly, however, both Mtr4 and Nop53 appear to occupy a broader region of the nucleolus than Rrp43 (Fig. 9A and B). Utp18, a SSU processome factor that binds pre-rRNA co-transcriptionally and recruits the exosome for the degradation of 5′-ETS after cleavage at A_0_^40^, shows a more compact nucleolar localization than Nop53 and Mrt4, and colocalized only partially with Rrp43 (Fig. 9C), confirming the observations described above of the exosome being concentrated in the same region as Rrp1, which might correspond to the Granular Component (GC). Because the exosome cofactors were more broadly present in the nucleolus than Rrp43, we compared their levels in the cells by immuno blot of TAP-tagged versions of these proteins, and show that these cofactors are indeed present in higher levels in the cells than Rrp43 (Fig. 9D), confirming the microscopy results, and in agreement with yeast protein abundance measurements^41^.

**Figure 9.**
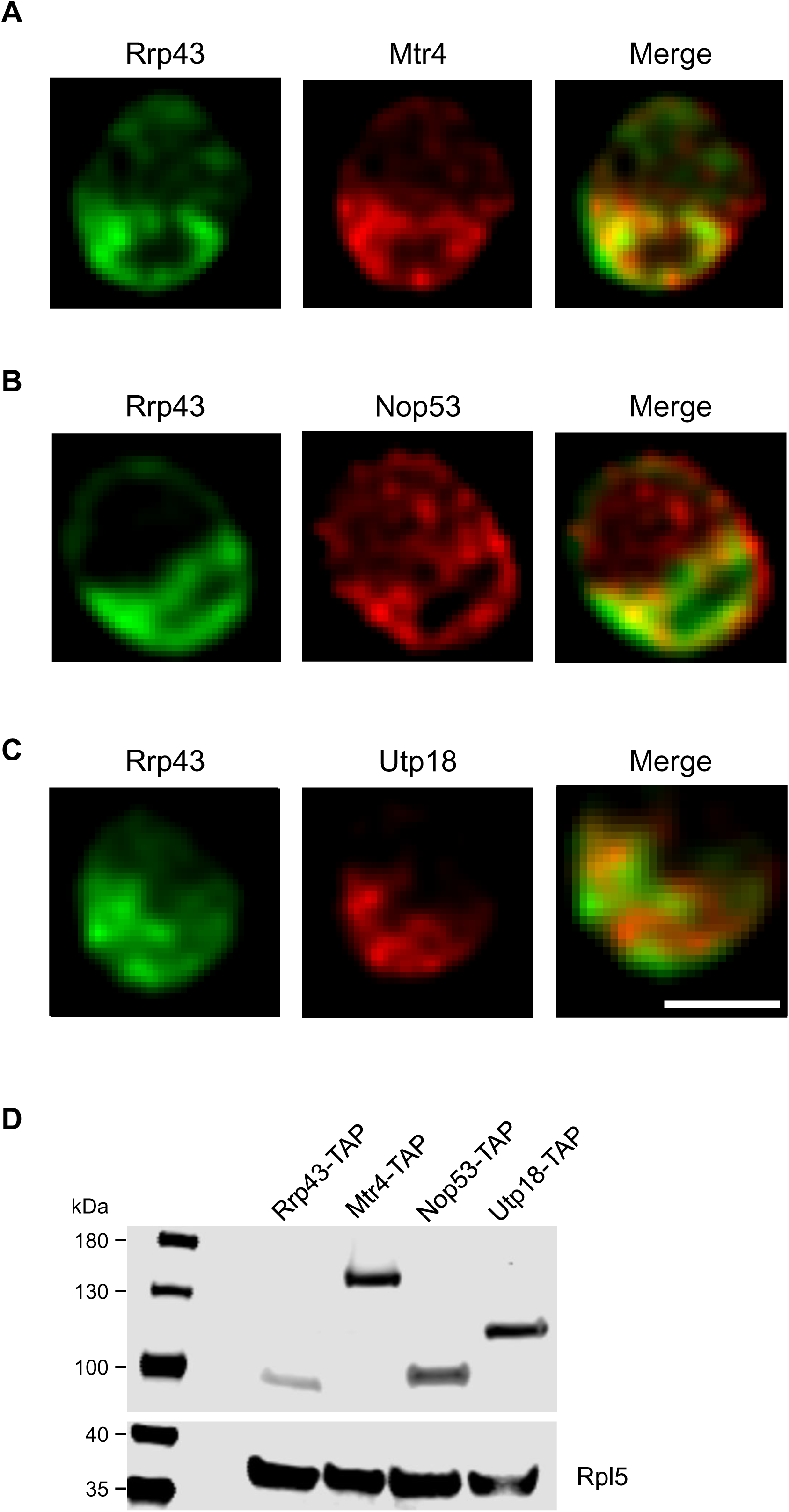
Rrp43 colocalizes with Mtr4 and Nop53 in the nucleolus but is less concentrated in the region containing Utp18. Images obtained by Z section of spinning disk confocal microscope of Rrp43-GFP compared to MTR4-mCherry (**A**), Nop53-mcherry (**B**) and UTP18-mCherry (**C**) in asynchronous cells. (**D**) Western blot showing the expression levels of the proteins analyzed, compared to ribosomal protein Rpl5, used as a loading control.

Also in accordance with the spinning disk confocal microscopy data described above, the use of image scanning microscopy (ISM) to analyze in more detail the localization of Rrp43 showed that it is not present at the rDNA, where Net1 is concentrated. Rather, Rrp43 is concentrated in a more external layer than that where Net1 is localized (Fig 10A). Furthermore, by comparing Rrp43 with Rpa190, it can be seen that they do not colocalize in the nucleolus and Rrp43 is in an outer layer relative to Rpa190 (Fig. 10B), which also confirms our previous data on Rrp44^27^. Rrp43 localization is also different from Nop56 (Fig. 10C), as also described above. Confirming the results from spinning disk confocal microscopy, Rrp43 colocalizes with Rrp1 in ISM (Fig. 10D). To exclude the possibility that these localization differences were due to the GFP fusion of Rrp43, a Rrp43-mCherry fusion was analyzed and compared to the rDNA binding protein Fob1 fused to GFP, and the results show that Rrp43 is outside of the region where Fob1 is concentrated (Fig. 10D), confirming that the tag does not influence Rrp43 localization, and that it is not in the Fibrillar Center (FC).

**Figure 10.**
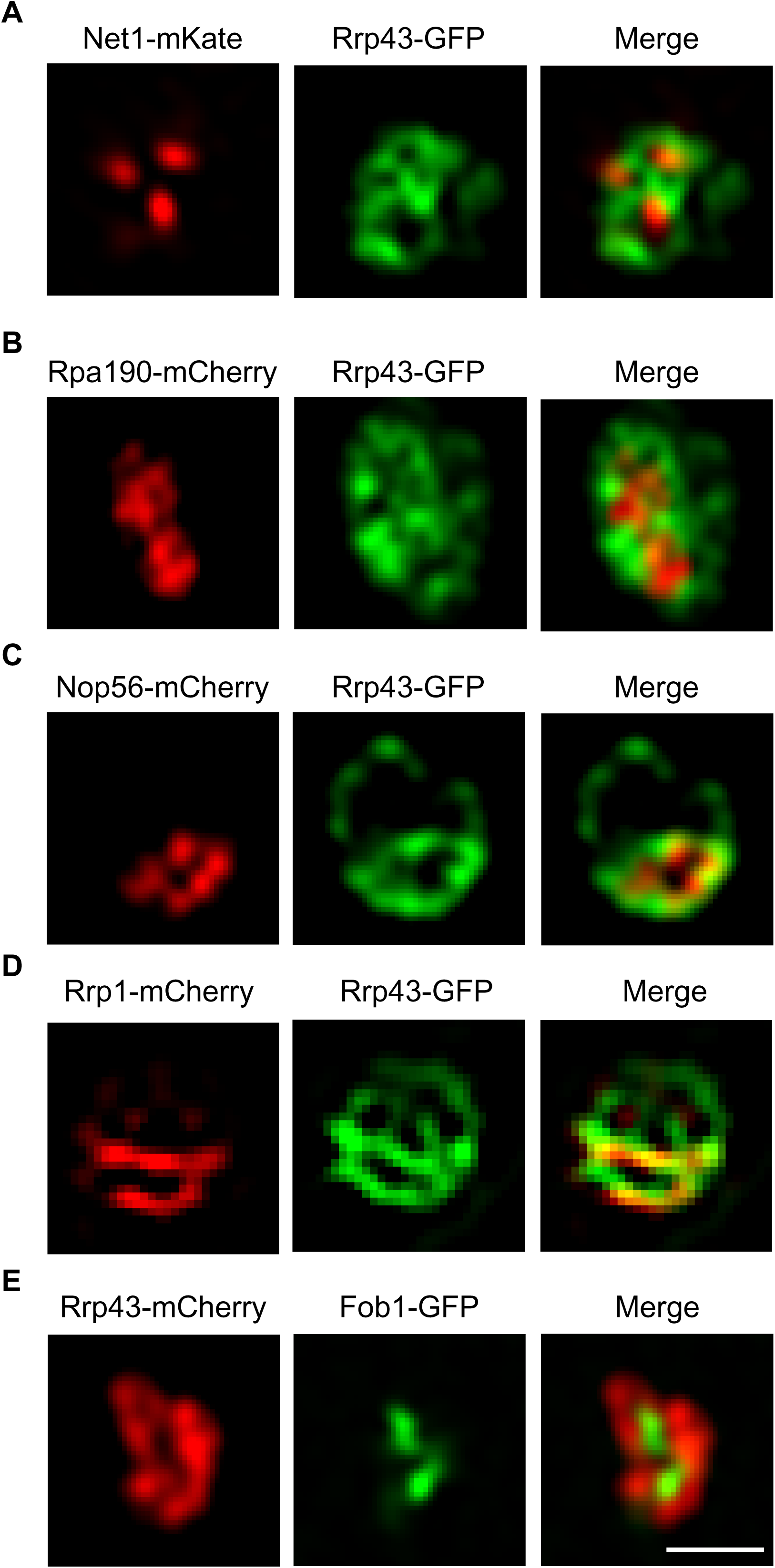
The concentration Rrp43 in nucleolus is lower in the rDNA region. Images obtained by image scanning high resolution microscope show the subcellular localization of Rrp43-GFP compared to Net1-mKate and Fob1-GFP (rDNA marker), Rpa190-mcherry and Nop56-mcherry (FC/DFC) in asynchronous cells. (**A**) Localization of Net1-mKate and Rrp43-GFP. (**B**) Rpa190-mcherry and Rrp43-GFP. (**C**) Nop56-mcherry and Rrp43-GFP. (**D**) Rrp1-mCherry and Rrp43-GFP. (**E**) Control Rrp43-mCherry and Fob1-GFP.

Combined, these results clearly show that despite being concentrated in the nucleolus, the exosome is not detected in the fibrillar center, where rDNA is concentrated and where Net1 and Fob1 localize, nor is the exosome enriched in the Dense Fibrillar Component (DFC), where Rpa190, Utp18 and Nop56 are concentrated, but rather the exosome is concentrated in the same region as Rrp1, Nop53 and Mtr4, which act at later pre-rRNA processing steps. Importantly, the last two are important for exosome recruitment to pre-60S rRNA.

### Correlative light and transmission electron microscopy (CLEM) reveals that exosome is concentrated in the granular component (GC) of the yeast nucleolus

Next, we aimed to identify the specific regions within the nucleolus in which the subunit of the exosome Rrp43 is concentrated. The nucleolus is most effectively studied using electron microscopy, which provides nanometer-resolution insights into the ultrastructural details of nuclear morphology^42–44^. Ultrastructure of the budding yeast nucleolus was initially categorized into three compartments based on their morphological appearance in transmission electron microscopy (TEM): the fibrillar center (FC), the dense fibrillar component (DFC), and the granular component (GC)^42,45^. CLEM analysis clearly shows that Rrp43-mCherry does not colocalize with the histone Hta1-GFP, which is concentrated in the nucleoplasm, whereas Rrp43 in the nucleolus (Fig. 11A), nor does Rrp43-neonGreen colocalizes with Net1-mKate, the latter bound to rDNA (Fig. 11B). When compared to Nop56-mCherry, Rrp43-neonGreen clearly localizes outside of the region where Nop56 is concentrated, which corresponds to the DFC (Fig. 11C). These results strongly suggested that the exosome is concentrated in the GC, where intermediate rRNA processing reactions take place.

**Figure 11.**
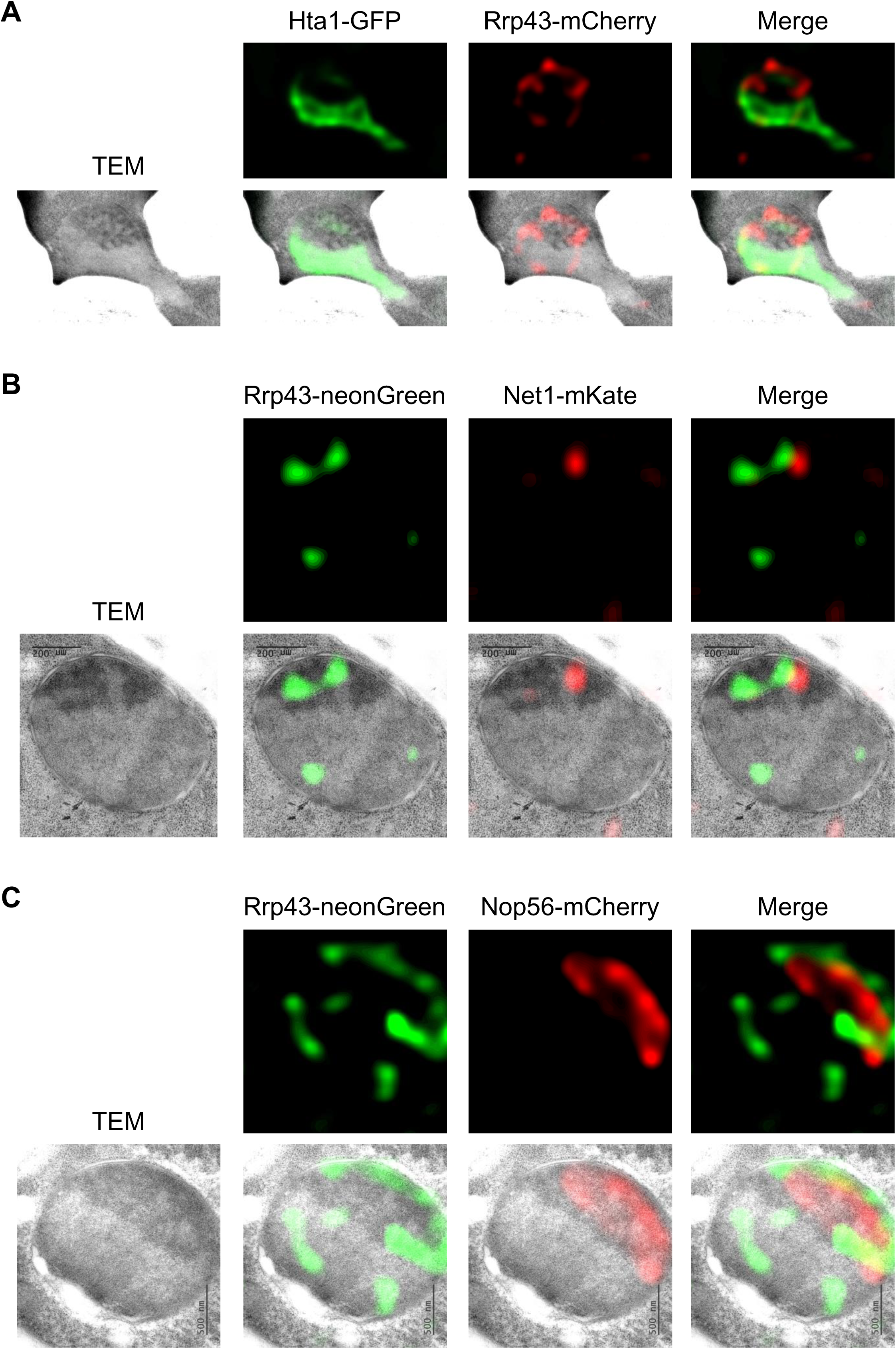
Correlative light and electron microscopy (CLEM) analysis of Rrp43 compared with nucleoplasmic (Hta1), fibrillar center (Net1), and dense fibrillar component (Nop56) nucleolar markers. (**A**) Overlay of TEM (transmission electron microscopy) and fluorescence microscopy images showing Rrp43-mCherry localization relative to Hta1-GFP fluorescence. (**B**) Overlay of TEM and fluorescence microscopy showing Rrp43-neonGreen localization relative to Net1-mKate fluorescence. (**C**) Overlay of TEM and fluorescence microscopy showing Rrp43-neonGreen localization relative to Nop56-mCherry fluorescence. A thickness section of approximately 150 to 180 nm was used for this analysis. Within this range, the resolution of pre-ribosomes in TEM (transmission electron microscopy) is reduced, and the fluorescence signals acquired are relative to this specific section thickness.

To further confirm our findings, three different fluorophores for tagging various nucleolar markers were used: Fob1-ypET (FC), Nop56-cyan (DFC), and Rrp43-mCherry. Additionally, Rrp1-mCherry was compared to Fob1-ypET and Nop56-TQ2. In exponential growth phase, we observed that Fob1 was surrounded by Nop56, and in turn, Nop56 was surrounded by Rrp43 or Rrp1 (Fig. 12), as predicted by the presence of three regions in the nucleolus. During rapamycin treatment, with the condensation of chromatin in the nucleolus, we noted the segregation of Fob1 (FC), Nop56 (DFC), and Rrp43/Rrp1 (GC), which continue to be concentrated in adjoining, but different nucleolar regions (Fig. 12). These results further confirm that three different regions are part of the yeast nucleolus, similar to human cells, and that the exosome is concentrated in the granular component.

**Figure 12.**
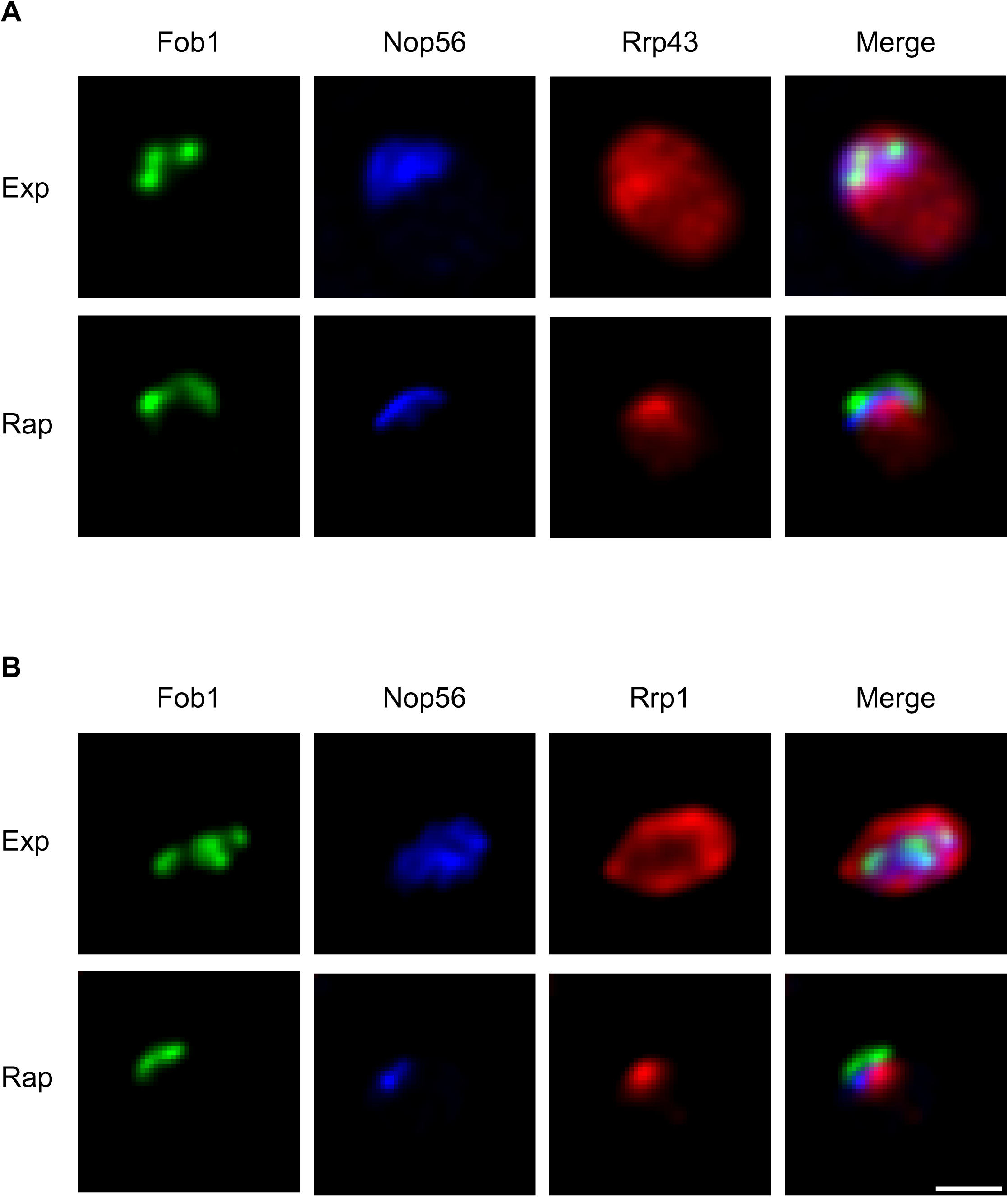
Segregation of proteins localized in GC, DFC, and FC regions of the nucleolus following transcriptional interruption. Z-section images acquired using a spinning disk confocal microscope show the localization of Rrp43-mCherry (**A**) and Rrp1-mCherry (**B**) in red, compared to Fob1-yPET (rDNA marker; green) or Nop56-TQ2 (DFC marker; blue). Localization was assessed in asynchronous cells during exponential growth and after rapamycin treatment.

## Discussion

The RNA exosome was first identified in *Saccharomyces cerevisiae* as an RNase involved in maturation and quality control of stable RNAs^7,8^. In the subsequent studies, it became clear that the exosome is a protein complex conserved throughout evolution, which is present in the nucleus and cytoplasm of eukaryotic cells, where it interacts with many cofactors and participates in different RNA processing and degradation pathways^17,46,47^.

In yeast, Exo10, composed of the exosome barrel and Rrp44, is present both in nucleus and cytoplasm, participating in different reactions in each of these subcellular compartments^48^. Exo11 has the additional nuclear-exclusive subunit Rrp6^49^. We have previously identified the nuclear import pathways of Rrp6 and of Rrp44 and shown that these exosome subunits have redundant mechanisms of transport to the nucleus, directly interacting with α-importin Srp1 and β-importins Kap95 and Sxm1^25,27^. Based on the evidences that Rrp6 and Rrp44 do not independently mediate the nuclear import of the exosome core^25,27^, we set out to investigate the transport pathway of the exosome subunits that form the complex core to the nucleus, where this complex participates in the essential process of ribosome maturation. With the exception of Rrp43, none of the nine exosome core subunits contain a nuclear localization signal (NLS) in the primary sequences. In the case of Rrp43, we identified a putative NLS in its primary sequence, but analysis of its structure in the exosome complex indicates that this NLS may not be readily accessible to karyopherins because of its partially internal localization in the Rrp43 tridimensional structure^21^. Despite the absence of a clearly exposed NLS, we show here that all exosome subunits localize mainly to the nucleus and are concentrated in the nucleolus, quantitatively confirming that the main function of the yeast exosome is the processing of pre-rRNAs. Its high concentration in the nucleoplasm also point to surveillance of nuclear RNAs as an important function of the exosome^7,47,50,51^.

One of the key questions regarding the nuclear transport of the exosome is whether it is transported as a whole core complex, or whether subcomplexes are transported independently and the exosome core is reassembled in the nucleus. Although the interactions between some of the exosome subunits are stronger, allowing the isolation of subcomplexes^52,53^, the presence of a putative NLS only in Rrp43 of the nine exosome barrel constituents might indicate that this subunit is important for the nuclear import of the complex as a whole. We tested this hypothesis by analyzing the effect of the depletion of Rrp43 on the subcellular localization of three other subunits, Rrp45, Rrp46, and Mtr3 (the last two interact directly with Rrp43) and show that the depletion of Rrp43 leads to a higher concentration of these three subunits in the cytoplasm. We hypothesize that in the absence of one core subunits the assembly and stability of the exosome complex is negatively affected, hindering its transport to the nucleus. Corroborating these findings, mutants of Rrp43 has been shown to destabilize the exosome complex^54^.

We have previously shown that Srp1 and Kap95 are the major karyopherins involved in the nuclear transport of Rrp6 and Rrp44^25,27^, and confirmed here their involvement in the transport of the other exosome subunits, with the surprising exception of Csl4 localization not being affected by the depletion of Kap95. This result may suggest that Csl4 can be transported by Srp1 associated with a β-karyopherin other than Kap95. Despite also being involved in the nuclear import of Rrp6, Sxm1 does not affect Csl4 localization, similar to the other two β-karyopherins tested, Kap123 and Msn5, which may indicate that Csl4, possibly bound to an NLS-containing factor, could be transported by the α-karyopherin Srp1 on its own, as has been described for other proteins^55,56^. Interestingly, Csl4 has been shown to associate less stably with the exosome core in protein co-purification experiments^53,57,58^, while associating strongly with Ski7^59^. Additionally, Arabidopsis exosome complex lacking Csl4 has been shown to remain nearly intact in size exclusion chromatography^58^. The observation shown here that Csl4 is not affected by Kap95 depletion, suggests that Csl4 might be transported to the nucleus independently, when not stably bound to the cytoplasmic exosome complex.

The SKI complex has been shown to recruit the exosome for cytoplasmic RNA degradation through the adaptor protein Ski7^31^, which binds the exosome through Csl4, at the top RNA binding ring, and Rrp43 and Mtr3, at the RNase PH ring^33,60^. Interestingly, Ski7 binding site on the exosome overlaps that of the nuclear exclusive subunit, Rrp6^21^, suggesting that Ski7 might play a role in the retention of the exosome core in the cytoplasm, hypothesis that was confirmed here by the observation that overexpression of Ski7 leads to higher concentration of Csl4, Rrp43 and Mtr3 in the cytoplasm. These results may indicate that the exosome can be split in subcomplexes under certain conditions. Accordingly, exosome subcomplexes have been purified from yeast cells through TAP chromatography followed by gel filtration^53^, in which a subcomplex containing the subunits Rrp4-Rrp41-Rrp42-Rrp45-Rrp44 was purified, almost exactly corresponding to the complement of the Csl4-Rrp43-Mtr3 subcomplex that was retained in the cytoplasm by the overexpression of Ski7.

The fact that the exosome core subunits do not contain identified NLS, however, raises the question of whether Rrp44, which is also part of the complex core and present in the cytoplasm, would be responsible for signaling the transport of the exosome to the nucleus after being bound by Srp1. As shown here, depletion of Rrp44 affects the nuclear concentration of RNA binding subunit Rrp40 and the RNase PH subunits Rrp43 and Mtr3, but not to the same extent as depletion of Rrp43 affects the localization of the RNase PH subunits Rrp45, Rrp46 and Mtr3. Combined, these results exclude the role of Rrp44 as the subunit signaling the exosome nuclear import and further confirm the importance of the stability of the exosome complex for its transport to the nucleus.

Strikingly, depletion of Rrp44, despite the little effect on exosome localization, affects the localization of RNA polymerase I subunit Rpa190. Although remaining concentrated in the nucleolus, Rpa190 spreads through the nucleoplasm in the absence of Rrp44. rRNA transcription effects on rRNA processing have already been described and denote the importance of co-transcriptional rRNA folding and interaction with r-proteins and ribosome assembly factors^30,61^. Rpa190 and other subunits of RNA pol I have been shown to interact with the transcription factors Spt4/Spt5, which interact physically and genetically with the Nrd1/Nab3/Sen1 complex and with the exosome, thereby facilitating the rRNA quality control^62^. The results shown here indicate that the cross-talk between these factors may result in the reverse effect of a defective exosome leading to a diffuse localization of RNA polymerase I. Interestingly, human exosome subunits have been reported to remain associated with perinucleolar bodies during mitosis, underlining the role of the exosome in nucleolar structure^63^. Furthermore, the recent identification of the human exosome subunits EXOSC7 (hRrp42) and EXOSC8 (hRrp43) in a complex with ZNF692, a protein that interacts with the nucleolar protein nucleophosmin 1 (NPM1), shows the multifunctional role of the exosome in ribosome formation and nucleolus arrangement^64^. NPM1 itself is a protein involved in a variety of cellular processes, including ribosomal maturation^65^.

Even though concentrated in the nucleolus, the exosome subunits do not exactly overlap Rpa190, suggesting a slightly different localization^27^. Here, we analyzed in more detail the subnucleolar localization of the exosome, comparing it with proteins known to associate with rDNA (Net1, Fob1)^36,66,67^, or that are part of the early (Nop56, Utp18)^68^, or of the late (Mtr4, Nop53)^69^ pre-rRNA processing steps. By analyzing the localization of these proteins by super-resolution microscopy, we were able to describe three different subnucleolar regions, one corresponding to rDNA, to which Net1 binds and can be considered the fibrillar center (FC), a second subcompartment, where Nop56 and Utp18 localize, which are components of the SSU processome, indicating the interface between FC and DFC (dense fibrillar component), and a third region, where Mtr4 and Nop53 are concentrated, proteins that participate at later stages of pre-rRNA processing, which would correspond to the granular component (GC). These results contradict earlier predictions that yeast would have only two nucleolar compartments^43^, but are in agreement with models predicting that the rRNA folding and change in the composition of pre-ribosomal particles are key for their transport from the nucleolus to the nucleoplasm, before reaching the cytoplasm, and with the presence of three compartments also in the yeast nucleolus^35,42,70^.

In summary, here we show that all exosome subunits are concentrated in the nucleolus, and that its NLS-containing catalytic subunits, Rrp6 and Rrp44, are not responsible for the nuclear import of the noncatalytic core of the complex. Importantly, we confirm that the yeast nucleolus is formed by three sub-compartments corresponding to the FC, DFC, and GC, and that the exosome is concentrated in the GC. Our analysis suggests a spatial segregation between rRNA synthesis and rRNA maturation and degradation by the exosome, underlining its participation in the late phases of pre-rRNA processing and quality control. Furthermore, we show for the first time that the potential competition between Rrp6 and Ski7 may play a role in the exosome core subcellular localization and consequently function control.

## Materials and Methods

### Construction of strains and yeast growth condition

Standard procedures were used for the propagation of yeast using YPD medium (1% yeast extract, 2% peptone and 2% glucose), or YNB medium (0.67% yeast nitrogen base, 0.5% (NH4)2SO4 and 2% glucose or galactose) supplemented with the required amino acids. Antibiotics Hygromycin (HPH - 200 μm/mL), Geneticin (G418 - 200 μm/mL) or Nourseothricin sulfate (cloNAT -100 μm/mL) were added to YPD medium at the indicated concentration.

Strains with genomic insertion of GFP or Neongreen (green), mCherry or mKate (red), yPET (yellow), Tq2 (blue), and TAP were constructed in two steps. A DNA fragment containing the fluorophore and selectable marker was PCR amplified using existing plasmid as template (Supplementary table 1) using specific primers (supplementary table 2). Next, yeast was transformed with the PCR product, allowing the targeting of the epitope at the 3’ end of genes by homologous recombination^51^. Alternatively, existing insertion can be used to exchange fluorophore using the epitope switching method^71^. Transformants were selected on either selective media for auxotrophy, or on antibiotic, selective marker, by PCR and microscopy.

### Fluorescence microscopy on living cells

After centrifugation, yeast cells were suspended in synthetic complete medium (DIFCO), mounted on a slide, and observed in the fluorescence microscope. Confocal microscopy was performed using a Nipkow-disk confocal system (Revolution; Andor) installed on a microscope (IX-81), featuring a confocal spinning disk unit (CSU22; Yokogawa) and a cooled electron multiplying charge-coupled device camera (DU 888; Andor). The system was controlled using the FAST mode of Revolution IQ software (Andor). Images were acquired using a 100× Plan Apo 1.4 NA oil immersion objective and a twofold lens in the optical path. Single laser lines used for excitation were diode-pumped solid-state lasers exciting GFP fluorescence at 488 nm (50 mW; Coherent) and mCherry fluorescence at 561 nm (50 mW; Cobolt jive), and a bi-bandpass emission filter (Em01-R488/568-15; Semrock) allowed collection of the green and red fluorescence. Pixel size was 65 nm. For quantification of nucleolar volume, z stacks of 40 images with a 250-nm z step were used. Exposure times varied from 0,1 to 1 s. Digital pictures were processed using ImageJ (National Institutes of Health, Bethesda, MD).

### Image analysis and quantification

Confocal images were imported into ImageJ software (https://imagej.nih.gov), and signal intensity measured from GFP tagged protein and their subnuclear localizations were analyzed using a dedicated image analysis pipeline (imageJ Macro). Cell border was determined based on transmission light in which cell wall is detected using a white line in the border. Nucleolar and nuclear segmentation was achieved using RNA polymerase I largest subunit Rpa190 tagged with mCherry, strongly enriched in the nucleolus, and detectable in the nucleoplasm. Cell was subdivided in nucleolus, nucleoplasm and cytoplasm, in which GFP signal was quantified relative to control cells bearing no GFP fusion. Images were analyzed using Z projection and Sum slice option.

Images were also analyzed in Z-section to compare with two different proteins that are in the same space in live cells. After acquisition, images were submitted to deconvolution treatment (Huygens Professional software) using relevant parameters: optical properties of our microscopic set-up were added, voxel size (x 65nm, y 65nm, Z 200nm), objectify numerical aperture (1.4), Refractive indexes of lens immersion oil (NA=1.517, embedding med oil 1.515). Deconvolution parameters used were: background estimation radius 0.7, relative background 0.0, Bleaching correction (if possible), brick mode (auto), PSFs per brick mode (off), PSFs per brick, manual mode (1), array detector reduction mode (auto). Image was saved as a channel in TIFF 16bit single file. Deconvolved images were imported into ImageJ software, Channel images were merged and analyzed.

### Image Scanning Microscopy acquisition and reconstruction

ISM was used to observe GFP and mCherry staining using an upgrade of the system described previously^72^. 2D images were acquired using an inverted microscope (TEi Nikon) equipped with a ×100 magnification, 1.49 NA objective (CFI SR APO 100XH ON 1.49 NIKON) and SCMOS camera (ORCA-Fusion, Hamamatsu). A commercial acquisition software (INSCOPER SA) enables to image multi-color 2D whole-cell. Fast diode lasers (Oxxius) with the wavelengths centered at 405nm, 488 nm (LBX-488-200-CSB) and 561nm were used to produce a TEM00 2.2-mm-diameter beam. The polarization beam was rotated with an angle of 5° before hitting a X4 Beam Expander beam (GBE04-A) and produced an 8.8 mm TEM00 beam. A fast-spatial light phase binary modulator (QXGA fourth dimensions) was conjugated to the image plane to create a specific pattern of illumination in one plane as in 200 streams. 2D image reconstruction was then performed as described previously and at GitHub (https://github.com/teamRIM/tutoRIM)^72,73^.

### Correlative Light and Electron microscopy of yeast nuclei

Culture of budding yeast in exponential phase were centrifuged for 1 minute at 3000 RPM, pellet of cells was cryofixed by high pressure freezing (EMPACT; Leica) and cryosubstituted with 0.1% uranyl acetate in acetone for 72h. Cells were embedded in a Lowicryl resin (HM-20) polymerized at −50°C. Sections of about 150-180 nm (purple color) were obtained in ultramicrotome (Reichert-Jung Ultracut E) and deposited in copper grids coated with Forvar. Grids were covered with a carbon layer of 5nm and quality of cryofixation was checked in 1,200× electron microscope (Jeol-JEM 1400)^74^.

For fluorescence acquisition, grids were deposited on microscope slides with citifluor® AF1 mounting solution and covered with a coverslip. A Nikon wide field microscopy for fluorescence was uused (NiKon TI-E/B inverted microscope and EMCCD camera (Ixon Ultra DU897-ANDOR). Lasers 470 (GFP excitation) and 640 (RFP excitation) were used in 100%, exposure time 0.5 ms to 2s, 0.2 µm step, 11 steps range 2 um (−1.0, +1.0).

After fluorescence imaging acquisition, slides were incubated for 1 hour into distilled water to separate slide and coverslip without damaging the grid and remove the mounting medium. Washed grids were contrasted using uranyless for 1 minute and Renolds lead citrate 3% for 2 min in NaOH saturated atmosphere. Jeol-JEM 1400 electron microscope was used to image cells on the grids using different zooms: one for the full cell, one for taking cells around and one for taking the nucleolus. Finally, images were aligned using ec-CLEM^75^. First deformation matrix was done with images of transmission electron microscopy (TEM) of cell and nucleolus. The second matrix was calculated between TEM and BF of fluorescence images using cell shape. The plugin of no rigid transformation matrix was used. Both matrixes are then applied to modify the fluorescence image. Modified fluorescence image and TEM image are then merged.

### Immunoblotting experiments

Protein samples were resolved by SDS-PAGE and transferred to nitrocellulose (NC) membrane (GE Healthcare). Membranes were incubated with primary antibodies against His-tag (Sigma-Aldrich), GFP (Sigma-Aldrich), or Pgk1 (Abcam) in phosphate-buffered saline (PBS)/Tween 20/nonfat milk. Secondary antibodies used were anti-rabbit (IRDye 680RD) or anti-mouse IgG (IRDye 800CW) conjugated to fluorophore (Licor). Western blots were developed using Odyssey^®^ Imaging Systems.

**Table 1.** List of Plasmids

| **Name** | **Selective marker, / purpose** | **Reference** |
| --- | --- | --- |
| pFA6-mCherry-HIS3 | HIS3, integrative / C-terminal tagging of endogenous genes with mCherry | <sup>76</sup> |
| pFA6a-mCherry-KIURA3 | URA3, integrative / C-terminal tagging of endogenous genes with mCherry | Gadal, unpublished |
| pFA6-GFP(S65T)-KIURA3 | URA3, integrative / C-terminal tagging of endogenous genes with GFP | <sup>71</sup> |
| pFA6-GFP-HIS | HIS, integrative / C-terminal tagging of endogenous genes with GFP | <sup>71</sup> |
| pFA6-GFP-KAN | KAN, integrative / C-terminal tagging of endogenous genes with GFP | <sup>71</sup> |
| YPET-URA | URA3 | Addgene |
| Tq2-URA | URA3 | Addgene |

**Table 2.**
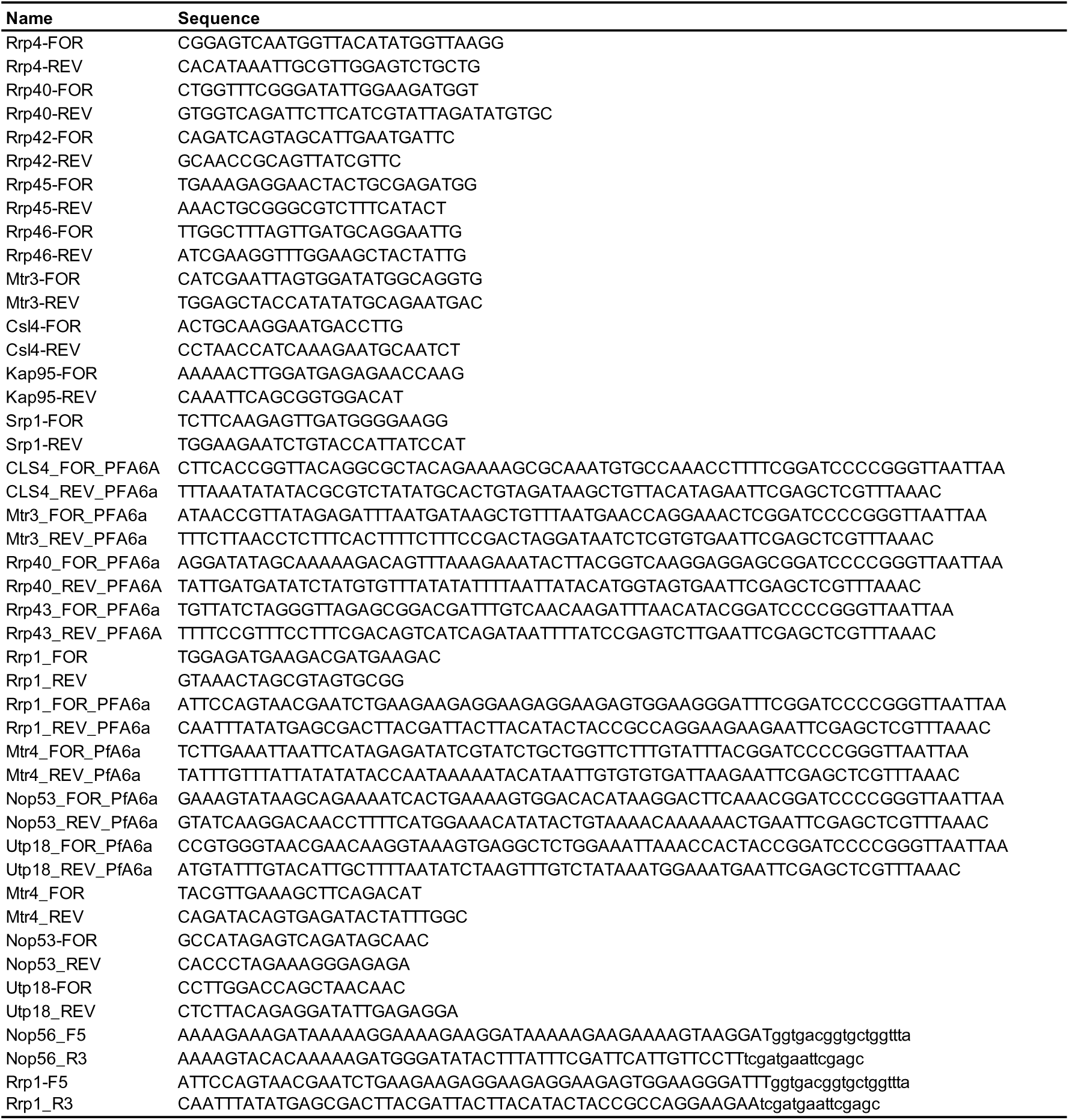
List of Primers

**Table 3.**
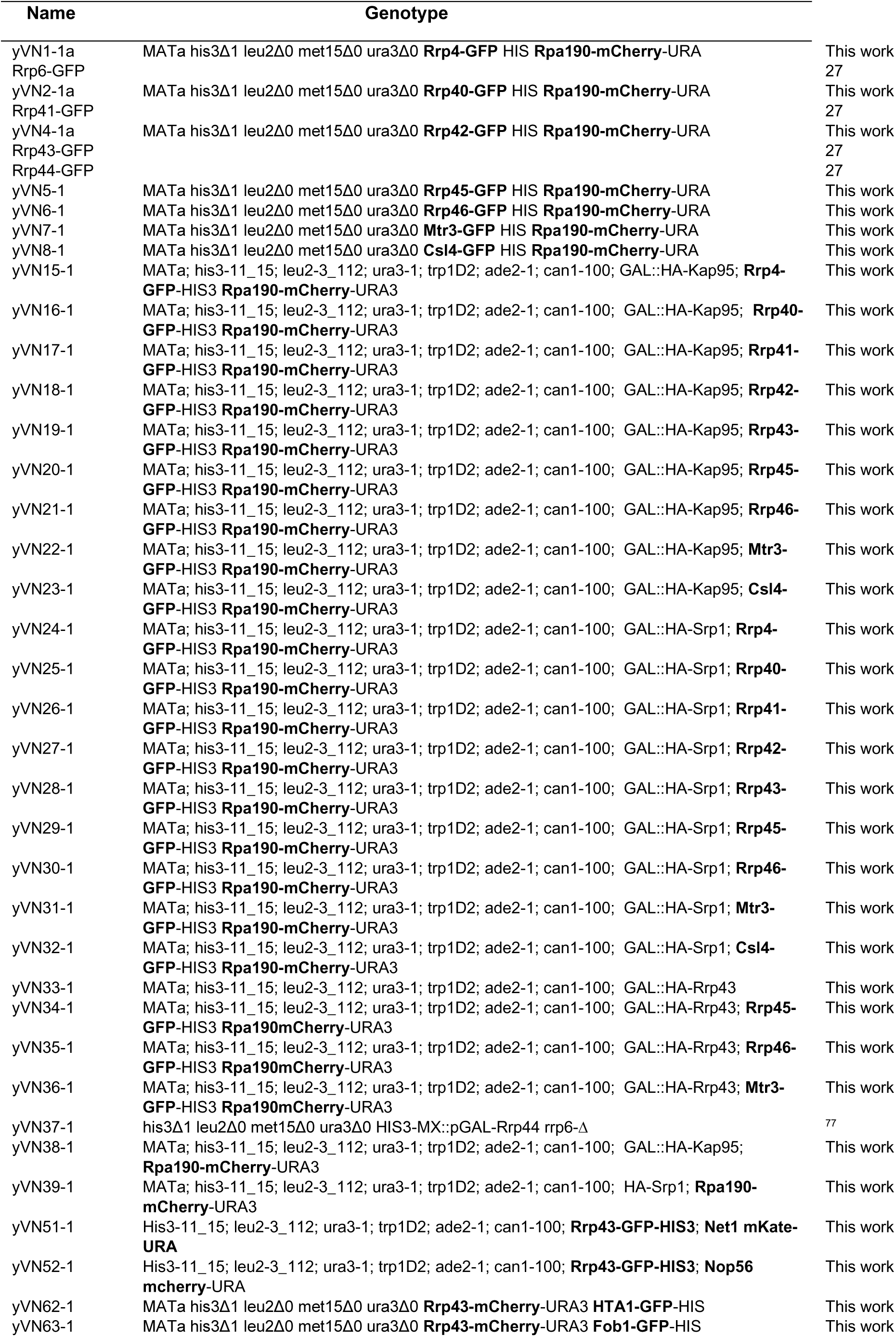
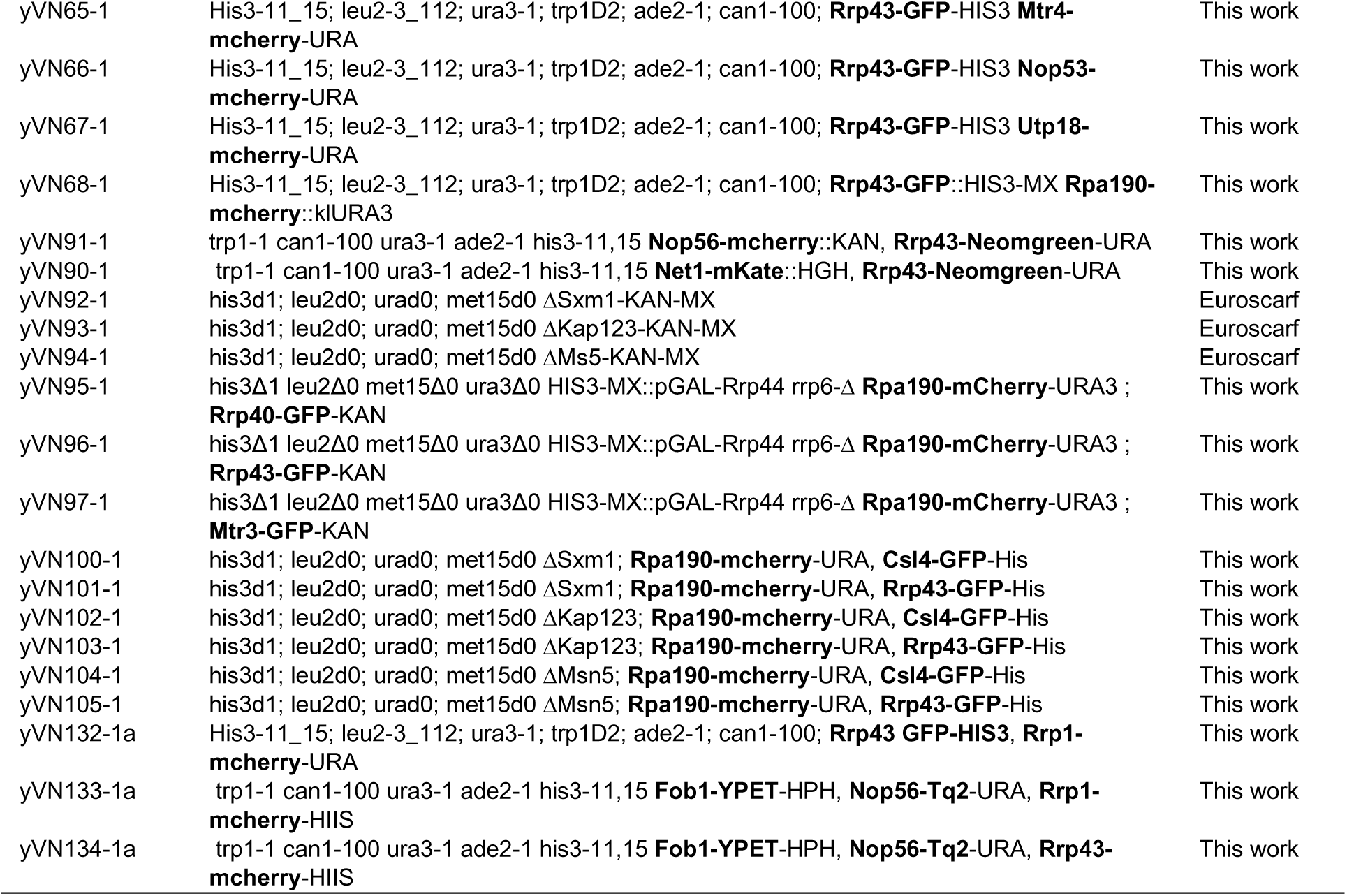
List of strains

## Acknowledgements

We are grateful to all members of the Oliveira and the Gadal laboratories for help, reagents, advice and discussion.

## Figures Legends

**Figure S1**

Schematics of the structure of the 35S pre-rRNA and major intermediates of the rRNA processing pathway in *S. cerevisiae*. The initial 35S pre-rRNA transcript, synthesized by RNA Polymerase I, contains sequences for mature 18S, 5.8S, and 25S rRNAs an internal and external transcribed spacer sequences (ITS/ETS). Processing intermediates formed during ribosome maturation are indicated. Exo- and endonucleolytic cleavage reactions are indicated. A “pacman” represents the exosome processing its intermediate substrates.

**Figure S2**

Strains expressing GFP-fused exosome subunits at endogenous levels grow as wild-type cells. Serial dilution of BY4741/Rpa190-mCherry strain expressing the exosome subunits fused to GFP in the endogenous loci, growing on YPD medium.

**Figure S3**

Amount of exosome subunits in different cell compartments. **(A**) nucleolus **(B)** nucleoplasm **(C)** cytoplasm. Amount calculated from Z-projection of high-resolution spinning-disk laser scanning confocal microscope images. Amounts were considered as (GFP signal of compartment) – (background of an equivalent area).

**Figure S4**

Area of nucleolus, nucleoplasm and cytoplasm in strains expressing GFP-tagged exosome subunits. (**A**) Cells areas were obtained from brightfield image. (**B**) Areas of nucleolus and (**C**) nucleoplasm were obtained from dual intensity of mCherry-tagged RNA polymerase I subunit Rpa190. (**D**) Cytoplasm area was obtained subtracting nucleus area from cell area.

**Figure S5**

Depletion of karyopherins Srp1 and Kap95. Strains *GAL1::HA-SRP1* and *GAL1::HA-KAP95* were grown in galactose medium, before being transferred to glucose medium for different periods of time. Strong depletion of both proteins can be observed after 6 hours in glucose (**A**), which coincides with the growth inhibition (**B**).

**Figure S6**

β-importins Kap123, Sxm1 and Msn5 are not involved in the nuclear import of the exosome subunits Csl4 and Rrp43. Csl4-GFP (**A**) and Rrp43-GFP (**B**) localization was analyzed in wild type or deletion strains for these three β-importins, showing that their deletion does not affect the nucleolar concentration of these core exosome subunits. RPA190-mCherry was used as a nucleolar marker. ImageJ plot is shown on the right, where green line represents GFP and red, RPA190-mCherry.

**Figure S7**

Depletion of Rrp43 and its effect on cell growth. Strains carrying the GAL1::HA-RRP43 construct were grown in galactose medium and subsequently transferred to glucose medium for varying durations. (**A**) Strong depletion of Rrp43 protein is observed after 6 hours in glucose medium. (**B**) Growth inhibition correlates with the depletion of Rrp43 protein. (**C**) Western blot detection of exosome core subunits Rrp45, Rrp46 and Mtr3 fused to GFP after depletion of Rrp43 show their decreased levels after 6 hours in glucose, when Rrp43 concentration drops significantly (**A**).

**Figure S8**

The absence of RNA exosome catalytic subunits leads to the mislocalization of RNA Pol I subunit Rpa190 and core RNA exosome subunits. (**A**) The localization of core subunits (Mtr3-GFP, Rrp43-GFP and Rrp40-GFP) was analyzed using Z section on a spinning disk confocal microscope. This analysis was performed in the strain *Δrrp6/GAL::RRP44/RPA190-mCherry* in galactose (presence of Rrp44) or in glucose medium for 6 hours (depletion of Rrp44). RPA190-mCherry was used as a nucleolar marker. (**B**) Western blot shows the depletion of HA-Rrp44 after growth in glucose-containing medium for different periods of time. Rpl5 was used as a loading control.

**Figure S9**

Ski7 does not affect the localization of the exosome catalytic subunits Rrp6 and Rrp44. (**A**) Localization of the plasmid-expressed GFP-fused exosome subunits in *Δski7/pMET15::SKI7-TAP/RPA190-mCherry* strain was analyzed by Z projection in spinning disk confocal microscope, showing that overexpression of Ski7 (-Met) does not affect these exosome subunits nuclear import. RPA190-mCherry was used as a nucleolar marker. (**B**) Csl4-GFP localization in wild-type or *Δski7/RPA190-mCherry* strain shows that endogenous levels of Ski7 do not affect Csl4 nucleolar concentration. (**C**) Western blot shows the effective Ski7 depletion in *Δski7/pMET15::SKI7-TAP/RPA190-mCherry* strain upon addition of methionine to the medium. Two different amounts of cell extracts were loaded on gel to show the higher expression levels of Ski7 upon induction of expression in the absence of methionine.

**Figure S10**

Localization of Fob1-yPET, Nop56-TQ2, and Rrp43/Rrp1-mCherry during exponential growth and after rapamycin treatment confirm that Rrp43 is concentrated outside the DFC in the nucleolus. Fluorescence microscopy images show the localization of Fob1-yPET (rDNA marker), Nop56-TQ2 (DFC marker), and Rrp43/Rrp1-mCherry in cells during exponential growth and following rapamycin treatment. Both raw images and deconvolved images are presented.

**Figure S11**

Same analysis as in Fig. 12, but with larger number of cells showing segregation of proteins localized in GC, DFC, and FC regions of the nucleolus following transcriptional interruption by the addition of rapamycin. Z-section images acquired using a spinning disk confocal microscope show the localization of Rrp43-mCherry (upper panels) and Rrp1-mCherry (lower panels) in red, compared to Fob1-yPET (rDNA marker; green) or Nop56-TQ2 (DFC marker; blue). Localization was assessed in asynchronous cells during exponential growth and after rapamycin treatment.

## Funding

This work was supported by a grant from Fundação de Amparo à Pesquisa do Estado de São Paulo (FAPESP - 20/00901-1 to C.C.O.), and by CBI (Centre de Biologie Integrative – Toulouse) to O.G. V.G.N. was supported by FAPESP fellowships (22/00071-4) and Research Internships Abroad (22/16740-2).

## Notes

### Competing Interest Statement

The authors have declared no competing interest.

## References

1. Scheer, U., and Rose, K.M. (1984). Localization of RNA polymerase I in interphase cells and mitotic chromosomes by light and electron microscopic immunocytochemistry. Proc Natl Acad Sci U S A 81, 1431–1435. 10.1073/pnas.81.5.1431.

2. Hugle, B., Hazan, R., Scheer, U., and Franke, W.W. (1985). Localization of ribosomal protein S1 in the granular component of the interphase nucleolus and its distribution during mitosis. J Cell Biol 100, 873–886. 10.1083/jcb.100.3.873.

3. Bassler, J., and Hurt, E. (2019). Eukaryotic Ribosome Assembly. Annu Rev Biochem 88, 281–306. 10.1146/annurev-biochem-013118-110817.

4. Chaker-Margot, M., Hunziker, M., Barandun, J., Dill, B.D., and Klinge, S. (2015). Stage-specific assembly events of the 6-MDa small-subunit processome initiate eukaryotic ribosome biogenesis. Nat Struct Mol Biol 22, 920–923. 10.1038/nsmb.3111.

5. Fatica, A., and Tollervey, D. (2002). Making ribosomes. Curr Opin Cell Biol 14, 313–318. 10.1016/s0955-0674(02)00336-8.

6. Kressler, D., Linder, P., and de La Cruz, J. (1999). Protein trans-acting factors involved in ribosome biogenesis in Saccharomyces cerevisiae. Mol Cell Biol 19, 7897–7912. 10.1128/MCB.19.12.7897.

7. Allmang, C., Kufel, J., Chanfreau, G., Mitchell, P., Petfalski, E., and Tollervey, D. (1999). Functions of the exosome in rRNA, snoRNA and snRNA synthesis. EMBO J 18, 5399–5410. 10.1093/emboj/18.19.5399.

8. Mitchell, P., Petfalski, E., Shevchenko, A., Mann, M., and Tollervey, D. (1997). The exosome: a conserved eukaryotic RNA processing complex containing multiple 3’-->5’ exoribonucleases. Cell 91, 457–466. 10.1016/s0092-8674(00)80432-8.

9. Henras, A.K., Plisson-Chastang, C., O’Donohue, M.F., Chakraborty, A., and Gleizes, P.E. (2015). An overview of pre-ribosomal RNA processing in eukaryotes. Wiley Interdiscip Rev RNA 6, 225–242. 10.1002/wrna.1269.

10. Zhang, L., Wu, C., Cai, G., Chen, S., and Ye, K. (2016). Stepwise and dynamic assembly of the earliest precursors of small ribosomal subunits in yeast. Genes Dev 30, 718–732. 10.1101/gad.274688.115.

11. Gasse, L., Flemming, D., and Hurt, E. (2015). Coordinated Ribosomal ITS2 RNA Processing by the Las1 Complex Integrating Endonuclease, Polynucleotide Kinase, and Exonuclease Activities. Mol Cell 60, 808–815. 10.1016/j.molcel.2015.10.021.

12. Konikkat, S., and Woolford, J.L., Jr. (2017). Principles of 60S ribosomal subunit assembly emerging from recent studies in yeast. Biochem J 474, 195–214. 10.1042/BCJ20160516.

13. Fromm, L., Falk, S., Flemming, D., Schuller, J.M., Thoms, M., Conti, E., and Hurt, E. (2017). Reconstitution of the complete pathway of ITS2 processing at the pre-ribosome. Nat Commun 8, 1787. 10.1038/s41467-017-01786-9.

14. Dez, C., Houseley, J., and Tollervey, D. (2006). Surveillance of nuclear-restricted pre-ribosomes within a subnucleolar region of Saccharomyces cerevisiae. EMBO J 25, 1534–1546. 10.1038/sj.emboj.7601035.

15. Choque, E., Schneider, C., Gadal, O., and Dez, C. (2018). Turnover of aberrant pre-40S pre-ribosomal particles is initiated by a novel endonucleolytic decay pathway. Nucleic Acids Res 46, 4699–4714. 10.1093/nar/gky116.

16. Darriere, T., Pilsl, M., Sarthou, M.K., Chauvier, A., Genty, T., Audibert, S., Dez, C., Leger-Silvestre, I., Normand, C., Henras, A.K., et al. (2019). Genetic analyses led to the discovery of a super-active mutant of the RNA polymerase I. PLoS Genet 15, e1008157. 10.1371/journal.pgen.1008157.

17. Januszyk, K., and Lima, C.D. (2014). The eukaryotic RNA exosome. Curr Opin Struct Biol 24, 132–140. 10.1016/j.sbi.2014.01.011.

18. Liu, Q., Greimann, J.C., and Lima, C.D. (2006). Reconstitution, activities, and structure of the eukaryotic RNA exosome. Cell 127, 1223–1237. 10.1016/j.cell.2006.10.037.

19. Dziembowski, A., Lorentzen, E., Conti, E., and Seraphin, B. (2007). A single subunit, Dis3, is essentially responsible for yeast exosome core activity. Nat Struct Mol Biol 14, 15–22. 10.1038/nsmb1184.

20. Lebreton, A., Tomecki, R., Dziembowski, A., and Seraphin, B. (2008). Endonucleolytic RNA cleavage by a eukaryotic exosome. Nature 456, 993–996. 10.1038/nature07480.

21. Makino, D.L., Schuch, B., Stegmann, E., Baumgartner, M., Basquin, C., and Conti, E. (2015). RNA degradation paths in a 12-subunit nuclear exosome complex. Nature 524, 54–58. 10.1038/nature14865.

22. Briggs, M.W., Burkard, K.T., and Butler, J.S. (1998). Rrp6p, the yeast homologue of the human PM-Scl 100-kDa autoantigen, is essential for efficient 5.8 S rRNA 3’ end formation. J Biol Chem 273, 13255–13263. 10.1074/jbc.273.21.13255.

23. Allmang, C., Petfalski, E., Podtelejnikov, A., Mann, M., Tollervey, D., and Mitchell, P. (1999). The yeast exosome and human PM-Scl are related complexes of 3’ --> 5’ exonucleases. Genes Dev 13, 2148–2158. 10.1101/gad.13.16.2148.

24. Burkard, K.T., and Butler, J.S. (2000). A nuclear 3’-5’ exonuclease involved in mRNA degradation interacts with Poly(A) polymerase and the hnRNA protein Npl3p. Mol Cell Biol 20, 604–616. 10.1128/MCB.20.2.604-616.2000.

25. Gonzales-Zubiate, F.A., Okuda, E.K., Da Cunha, J.P.C., and Oliveira, C.C. (2017). Identification of karyopherins involved in the nuclear import of RNA exosome subunit Rrp6 in Saccharomyces cerevisiae. J Biol Chem 292, 12267–12284. 10.1074/jbc.M116.772376.

26. Jin, R., Dobry, C.J., McCown, P.J., and Kumar, A. (2008). Large-scale analysis of yeast filamentous growth by systematic gene disruption and overexpression. Mol Biol Cell 19, 284–296. 10.1091/mbc.e07-05-0519.

27. Okuda, E.K., Gonzales-Zubiate, F.A., Gadal, O., and Oliveira, C.C. (2020). Nucleolar localization of the yeast RNA exosome subunit Rrp44 hints at early pre-rRNA processing as its main function. J Biol Chem 295, 11195–11213. 10.1074/jbc.RA120.013589.

28. Lau, B., Cheng, J.D., Flemming, D., La Venuta, G., Berninghausen, O., Beckmann, R., and Hurt, E. (2021). Structure of the Maturing 90S Pre-ribosome in Association with the RNA Exosome. Molecular Cell 81, 293-+. 10.1016/j.molcel.2020.11.009.

29. Uchida, M., Sun, Y., McDermott, G., Knoechel, C., Le Gros, M.A., Parkinson, D., Drubin, D.G., and Larabell, C.A. (2011). Quantitative analysis of yeast internal architecture using soft X-ray tomography. Yeast 28, 227–236. 10.1002/yea.1834.

30. Normand, C., Dez, C., Dauban, L., Queille, S., Danche, S., Abderrahmane, S., Beckouet, F., and Gadal, O. (2024). RNA polymerase I mutant affects ribosomal RNA processing and ribosomal DNA stability. RNA Biol 21, 1–16. 10.1080/15476286.2024.2381910.

31. van Hoof, A., Staples, R.R., Baker, R.E., and Parker, R. (2000). Function of the ski4p (Csl4p) and Ski7p proteins in 3’-to-5’ degradation of mRNA. Mol Cell Biol 20, 8230–8243. 10.1128/MCB.20.21.8230-8243.2000.

32. Tomecki, R., Drazkowska, K., Kobylecki, K., and Tudek, A. (2023). SKI complex: A multifaceted cytoplasmic RNA exosome cofactor in mRNA metabolism with links to disease, developmental processes, and antiviral responses. Wiley Interdiscip Rev RNA 14, e1795. 10.1002/wrna.1795.

33. Keidel, A., Kogel, A., Reichelt, P., Kowalinski, E., Schafer, I.B., and Conti, E. (2023). Concerted structural rearrangements enable RNA channeling into the cytoplasmic Ski238-Ski7-exosome assembly. Mol Cell 83, 4093–4105 e4097. 10.1016/j.molcel.2023.09.037.

34. Wasmuth, E.V., Januszyk, K., and Lima, C.D. (2014). Structure of an Rrp6-RNA exosome complex bound to poly(A) RNA. Nature 511, 435–439. 10.1038/nature13406.

35. Tartakoff, A.M., Chen, L., Raghavachari, S., Gitiforooz, D., Dhinakaran, A., Ni, C.L., Pasadyn, C., Mahabeleshwar, G.H., Pasadyn, V., and Woolford, J.L., Jr. (2021). The nucleolus as a polarized coaxial cable in which the rDNA axis is surrounded by dynamic subunit-specific phases. Curr Biol 31, 2507–2519 e2504. 10.1016/j.cub.2021.03.041.

36. Hannig, K., Babl, V., Hergert, K., Maier, A., Pilsl, M., Schachner, C., Stockl, U., Milkereit, P., Tschochner, H., Seufert, W., and Griesenbeck, J. (2019). The C-terminal region of Net1 is an activator of RNA polymerase I transcription with conserved features from yeast to human. PLoS Genet 15, e1008006. 10.1371/journal.pgen.1008006.

37. Dominique, C., Maiga, N.K., Mendez-Godoy, A., Pillet, B., Hamze, H., Leger-Silvestre, I., Henry, Y., Marchand, V., Gomes Neto, V., Dez, C., et al. (2024). The dual life of disordered lysine-rich domains of snoRNPs in rRNA modification and nucleolar compaction. Nat Commun 15, 9415. 10.1038/s41467-024-53805-1.

38. Tsang, C.K., Bertram, P.G., Ai, W., Drenan, R., and Zheng, X.F. (2003). Chromatin-mediated regulation of nucleolar structure and RNA Pol I localization by TOR. EMBO J 22, 6045–6056. 10.1093/emboj/cdg578.

39. Horsey, E.W., Jakovljevic, J., Miles, T.D., Harnpicharnchai, P., and Woolford, J.L., Jr. (2004). Role of the yeast Rrp1 protein in the dynamics of pre-ribosome maturation. RNA 10, 813–827. 10.1261/rna.5255804.

40. Thoms, M., Thomson, E., Bassler, J., Gnadig, M., Griesel, S., and Hurt, E. (2015). The Exosome Is Recruited to RNA Substrates through Specific Adaptor Proteins. Cell 162, 1029–1038. 10.1016/j.cell.2015.07.060.

41. Ho, B., Baryshnikova, A., and Brown, G.W. (2018). Unification of Protein Abundance Datasets Yields a Quantitative Saccharomyces cerevisiae Proteome. Cell Syst 6, 192–205 e193. 10.1016/j.cels.2017.12.004.

42. Leger-Silvestre, I., Trumtel, S., Noaillac-Depeyre, J., and Gas, N. (1999). Functional compartmentalization of the nucleus in the budding yeast Saccharomyces cerevisiae. Chromosoma 108, 103–113. 10.1007/s004120050357.

43. Thelen, N., Defourny, J., Lafontaine, D.L.J., and Thiry, M. (2021). Visualization of Chromatin in the Yeast Nucleus and Nucleolus Using Hyperosmotic Shock. Int J Mol Sci 22. 10.3390/ijms22031132.

44. Thiry, M., and Lafontaine, D.L. (2005). Birth of a nucleolus: the evolution of nucleolar compartments. Trends Cell Biol 15, 194–199. 10.1016/j.tcb.2005.02.007.

45. Leger-Silvestre, I., Noaillac-Depeyre, J., Faubladier, M., and Gas, N. (1997). Structural and functional analysis of the nucleolus of the fission yeast Schizosaccharomyces pombe. Eur J Cell Biol 72, 13–23.

46. Aloy, P., Ciccarelli, F.D., Leutwein, C., Gavin, A.C., Superti-Furga, G., Bork, P., Bottcher, B., and Russell, R.B. (2002). A complex prediction: three-dimensional model of the yeast exosome. EMBO Rep 3, 628–635. 10.1093/embo-reports/kvf135.

47. Kilchert, C., Wittmann, S., and Vasiljeva, L. (2016). The regulation and functions of the nuclear RNA exosome complex. Nat Rev Mol Cell Biol 17, 227–239. 10.1038/nrm.2015.15.

48. Houseley, J., LaCava, J., and Tollervey, D. (2006). RNA-quality control by the exosome. Nat Rev Mol Cell Biol 7, 529–539. 10.1038/nrm1964.

49. Wasmuth, E.V., and Lima, C.D. (2012). Exo- and endoribonucleolytic activities of yeast cytoplasmic and nuclear RNA exosomes are dependent on the noncatalytic core and central channel. Mol Cell 48, 133–144. 10.1016/j.molcel.2012.07.012.

50. Schneider, C., and Tollervey, D. (2013). Threading the barrel of the RNA exosome. Trends Biochem Sci 38, 485–493. 10.1016/j.tibs.2013.06.013.

51. Ghaemmaghami, S., Huh, W.K., Bower, K., Howson, R.W., Belle, A., Dephoure, N., O’Shea, E.K., and Weissman, J.S. (2003). Global analysis of protein expression in yeast. Nature 425, 737–741. 10.1038/nature02046.

52. Oliveira, C.C., Gonzales, F.A., and Zanchin, N.I. (2002). Temperature-sensitive mutants of the exosome subunit Rrp43p show a deficiency in mRNA degradation and no longer interact with the exosome. Nucleic Acids Res 30, 4186–4198. 10.1093/nar/gkf545.

53. Hernandez, H., Dziembowski, A., Taverner, T., Seraphin, B., and Robinson, C.V. (2006). Subunit architecture of multimeric complexes isolated directly from cells. EMBO Rep 7, 605–610. 10.1038/sj.embor.7400702.

54. Lourenco, R.F., Leme, A.F., and Oliveira, C.C. (2013). Proteomic analysis of yeast mutant RNA exosome complexes. J Proteome Res 12, 5912–5922. 10.1021/pr400972x.

55. Nitahara-Kasahara, Y., Kamata, M., Yamamoto, T., Zhang, X., Miyamoto, Y., Muneta, K., Iijima, S., Yoneda, Y., Tsunetsugu-Yokota, Y., and Aida, Y. (2007). Novel nuclear import of Vpr promoted by importin alpha is crucial for human immunodeficiency virus type 1 replication in macrophages. J Virol 81, 5284–5293. 10.1128/JVI.01928-06.

56. Miyamoto, Y., Hieda, M., Harreman, M.T., Fukumoto, M., Saiwaki, T., Hodel, A.E., Corbett, A.H., and Yoneda, Y. (2002). Importin alpha can migrate into the nucleus in an importin beta- and Ran-independent manner. EMBO J 21, 5833–5842. 10.1093/emboj/cdf569.

57. Synowsky, S.A., van Wijk, M., Raijmakers, R., and Heck, A.J. (2009). Comparative multiplexed mass spectrometric analyses of endogenously expressed yeast nuclear and cytoplasmic exosomes. J Mol Biol 385, 1300–1313. 10.1016/j.jmb.2008.11.011.

58. Chekanova, J.A., Gregory, B.D., Reverdatto, S.V., Chen, H., Kumar, R., Hooker, T., Yazaki, J., Li, P., Skiba, N., Peng, Q., et al. (2007). Genome-wide high-resolution mapping of exosome substrates reveals hidden features in the Arabidopsis transcriptome. Cell 131, 1340–1353. 10.1016/j.cell.2007.10.056.

59. van Hoof, A., Frischmeyer, P.A., Dietz, H.C., and Parker, R. (2002). Exosome-mediated recognition and degradation of mRNAs lacking a termination codon. Science 295, 2262–2264. 10.1126/science.1067272.

60. Kowalinski, E., Kogel, A., Ebert, J., Reichelt, P., Stegmann, E., Habermann, B., and Conti, E. (2016). Structure of a Cytoplasmic 11-Subunit RNA Exosome Complex. Mol Cell 63, 125–134. 10.1016/j.molcel.2016.05.028.

61. Huffines, A.K., Engel, K.L., French, S.L., Zhang, Y., Viktorovskaya, O.V., and Schneider, D.A. (2022). Rate of transcription elongation and sequence-specific pausing by RNA polymerase I directly influence rRNA processing. J Biol Chem 298, 102730. 10.1016/j.jbc.2022.102730.

62. Lepore, N., and Lafontaine, D.L. (2011). A functional interface at the rDNA connects rRNA synthesis, pre-rRNA processing and nucleolar surveillance in budding yeast. PLoS One 6, e24962. 10.1371/journal.pone.0024962.

63. Fomproix, N., and Hernandez-Verdun, D. (1999). Effects of anti-PM-Scl 100 (Rrp6p exonuclease) antibodies on prenucleolar body dynamics at the end of mitosis. Exp Cell Res 251, 452–464. 10.1006/excr.1999.4578.

64. Lafita-Navarro, M.C., Hao, Y.H., Jiang, C., Jang, S., Chang, T.C., Brown, I.N., Venkateswaran, N., Maurais, E., Stachera, W., Zhang, Y., et al. (2023). ZNF692 organizes a hub specialized in 40S ribosomal subunit maturation enhancing translation in rapidly proliferating cells. Cell Rep 42, 113280. 10.1016/j.celrep.2023.113280.

65. Box, J.K., Paquet, N., Adams, M.N., Boucher, D., Bolderson, E., O’Byrne, K.J., and Richard, D.J. (2016). Nucleophosmin: from structure and function to disease development. BMC Mol Biol 17, 19. 10.1186/s12867-016-0073-9.

66. Shou, W., Sakamoto, K.M., Keener, J., Morimoto, K.W., Traverso, E.E., Azzam, R., Hoppe, G.J., Feldman, R.M., DeModena, J., Moazed, D., et al. (2001). Net1 stimulates RNA polymerase I transcription and regulates nucleolar structure independently of controlling mitotic exit. Mol Cell 8, 45–55. 10.1016/s1097-2765(01)00291-x.

67. Wai, H., Johzuka, K., Vu, L., Eliason, K., Kobayashi, T., Horiuchi, T., and Nomura, M. (2001). Yeast RNA polymerase I enhancer is dispensable for transcription of the chromosomal rRNA gene and cell growth, and its apparent transcription enhancement from ectopic promoters requires Fob1 protein. Mol Cell Biol 21, 5541–5553. 10.1128/MCB.21.16.5541-5553.2001.

68. Chaker-Margot, M., Barandun, J., Hunziker, M., and Klinge, S. (2017). Architecture of the yeast small subunit processome. Science 355. 10.1126/science.aal1880.

69. Bagatelli, F.F.M., de Luna Vitorino, F.N., da Cunha, J.P.C., and Oliveira, C.C. (2021). The ribosome assembly factor Nop53 has a structural role in the formation of nuclear pre-60S intermediates, affecting late maturation events. Nucleic Acids Res 49, 7053–7074. 10.1093/nar/gkab494.

70. LaPeruta, A.J., Micic, J., and Woolford, J.L., Jr. (2023). Additional principles that govern the release of pre-ribosomes from the nucleolus into the nucleoplasm in yeast. Nucleic Acids Res 51, 10867–10883. 10.1093/nar/gkac430.

71. Sung, M.K., Ha, C.W., and Huh, W.K. (2008). A vector system for efficient and economical switching of C-terminal epitope tags in Saccharomyces cerevisiae. Yeast 25, 301–311. 10.1002/yea.1588.

72. Mangeat, T., Labouesse, S., Allain, M., Negash, A., Martin, E., Guenole, A., Poincloux, R., Estibal, C., Bouissou, A., Cantaloube, S., et al. (2021). Super-resolved live-cell imaging using random illumination microscopy. Cell Rep Methods 1, 100009. 10.1016/j.crmeth.2021.100009.

73. Arnould, C., Rocher, V., Saur, F., Bader, A.S., Muzzopappa, F., Collins, S., Lesage, E., Le Bozec, B., Puget, N., Clouaire, T., et al. (2023). Chromatin compartmentalization regulates the response to DNA damage. Nature 623, 183–192. 10.1038/s41586-023-06635-y.

74. Normand, C., Berthaud, M., Gadal, O., and Leger-Silvestre, I. (2016). Correlative Light and Electron Microscopy of Nucleolar Transcription in Saccharomyces cerevisiae. Methods Mol Biol 1455, 29–40. 10.1007/978-1-4939-3792-9_3.

75. Paul-Gilloteaux, P., Heiligenstein, X., Belle, M., Domart, M.C., Larijani, B., Collinson, L., Raposo, G., and Salamero, J. (2017). eC-CLEM: flexible multidimensional registration software for correlative microscopies. Nat Methods 14, 102–103. 10.1038/nmeth.4170.

76. Bachellier-Bassi, S., Gadal, O., Bourout, G., and Nehrbass, U. (2008). Cell cycle-dependent kinetochore localization of condensin complex in. J Struct Biol 162, 248–259. 10.1016/j.jsb.2008.01.002.

77. Schneider, C., Leung, E., Brown, J., and Tollervey, D. (2009). The N-terminal PIN domain of the exosome subunit Rrp44 harbors endonuclease activity and tethers Rrp44 to the yeast core exosome. Nucleic Acids Res 37, 1127–1140. 10.1093/nar/gkn1020.

