## Supplementary figures and images for "New Insights into nuclear import and nucleolar localization of yeast RNA exosome subunits"

### Supplemental figures

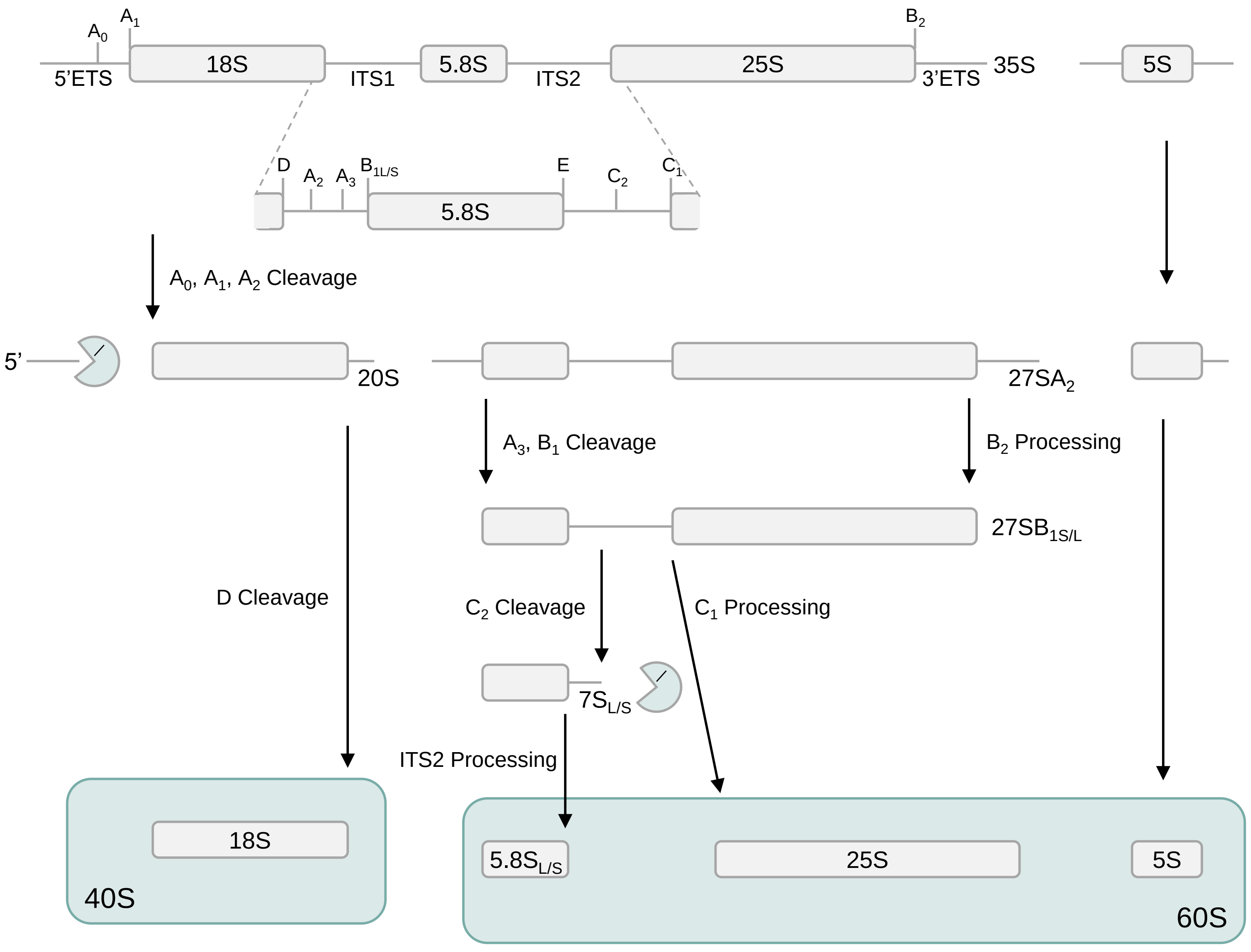

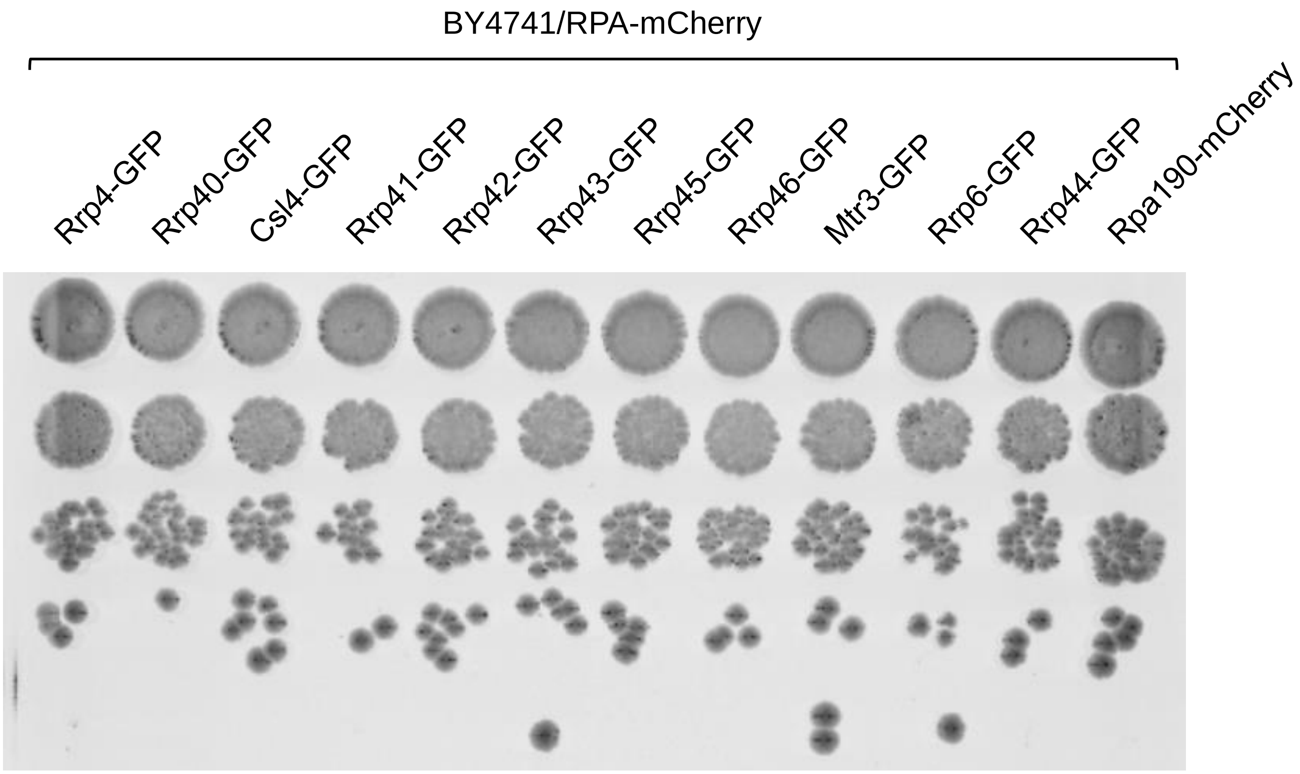

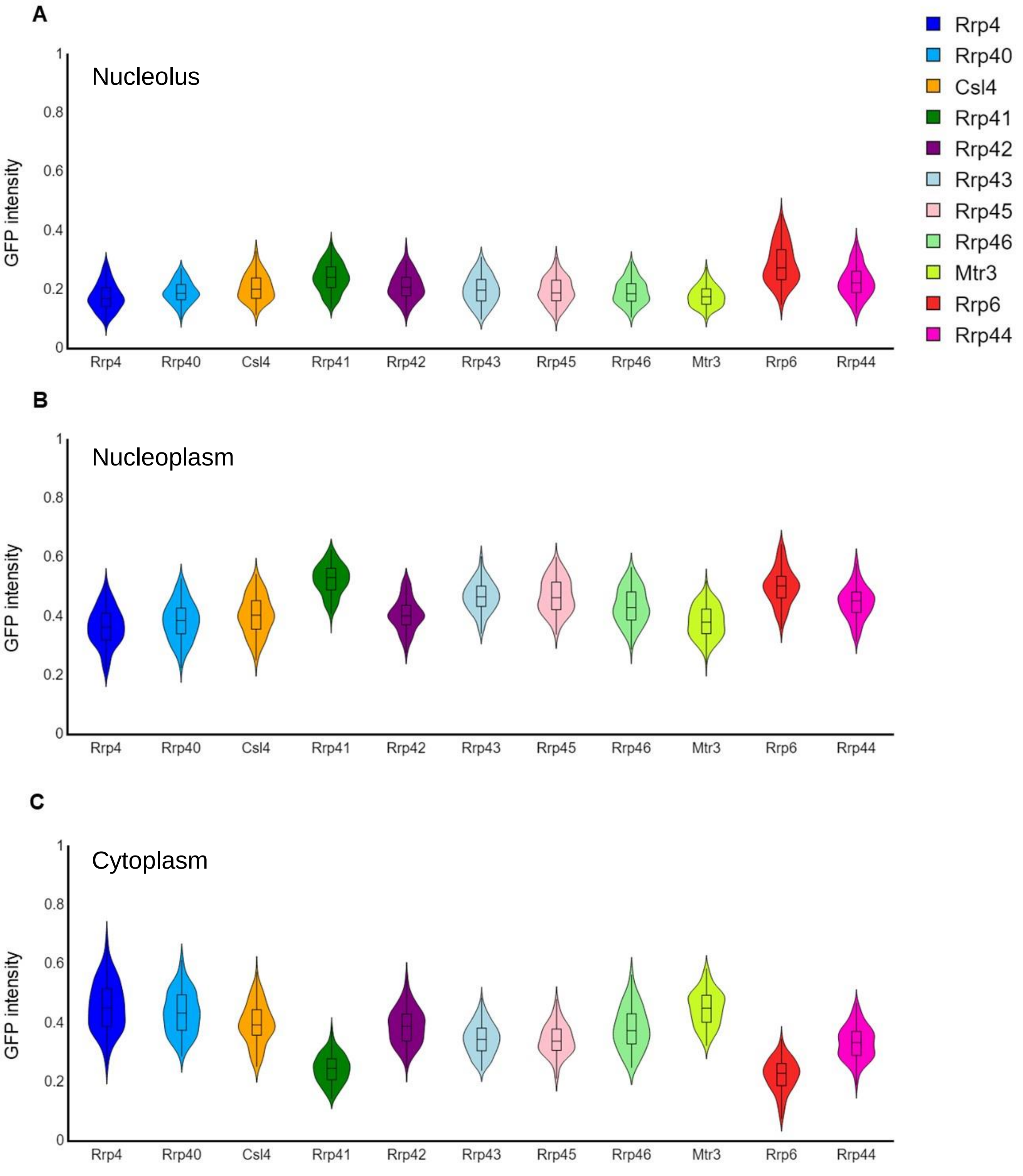

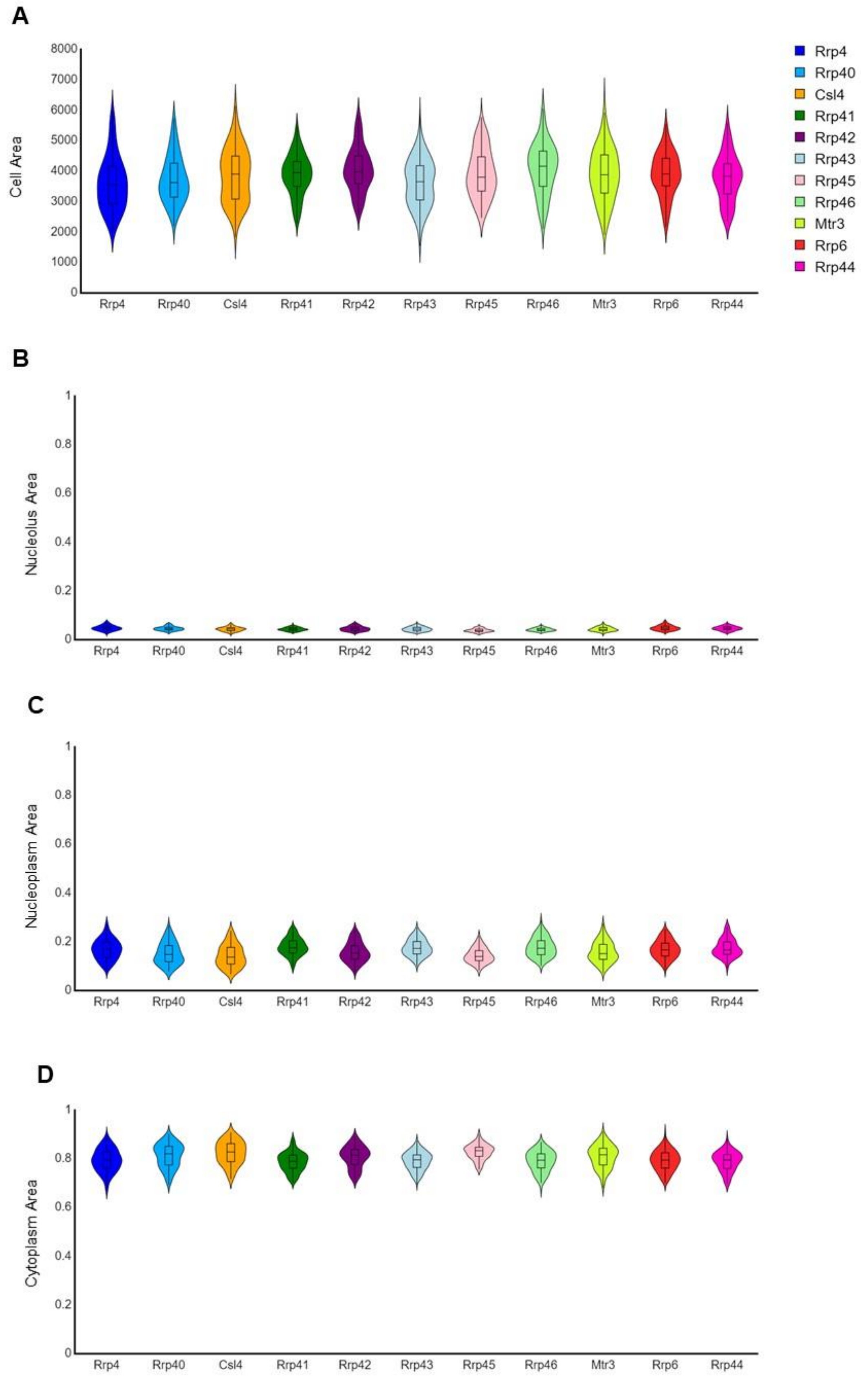

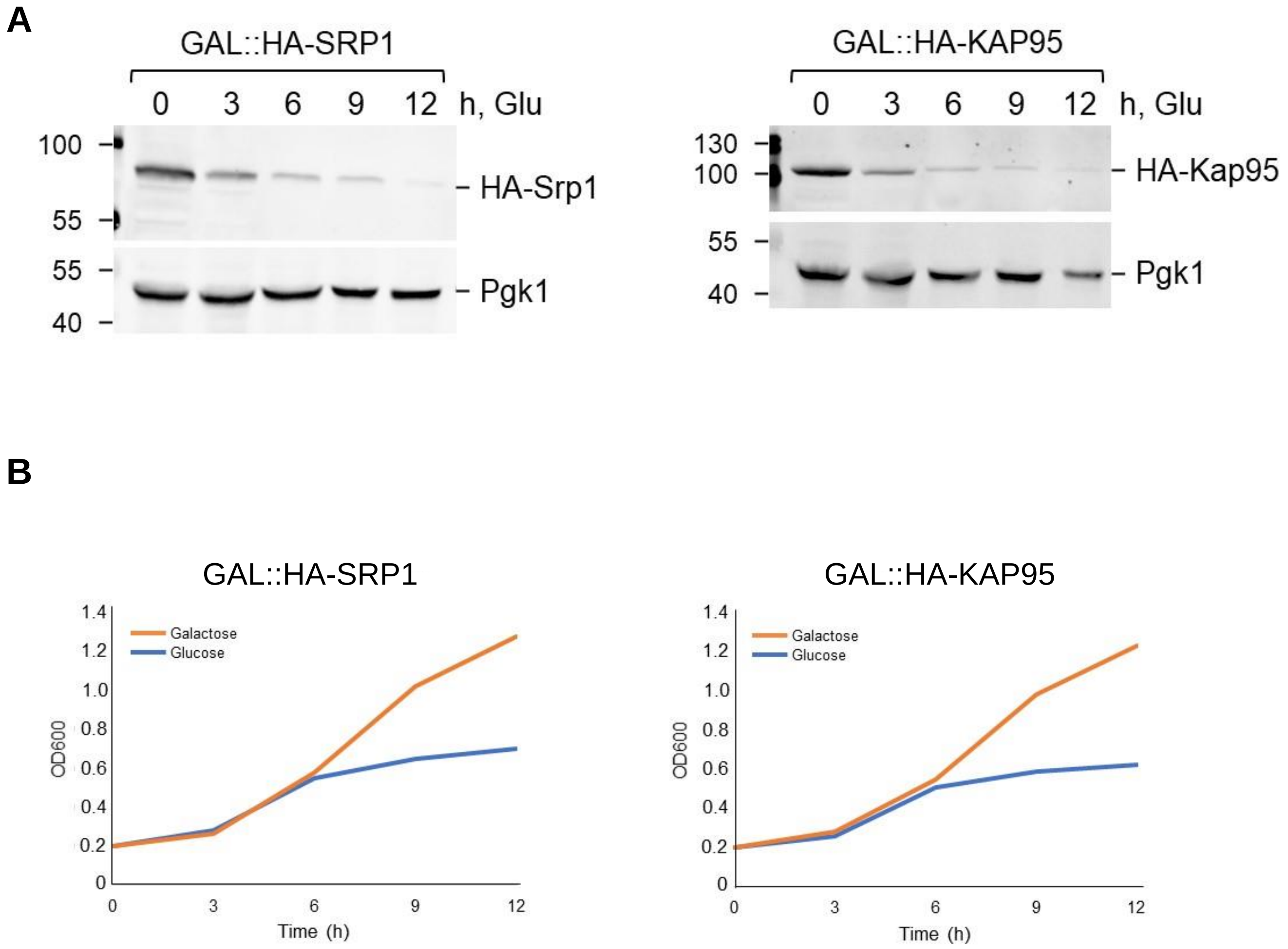

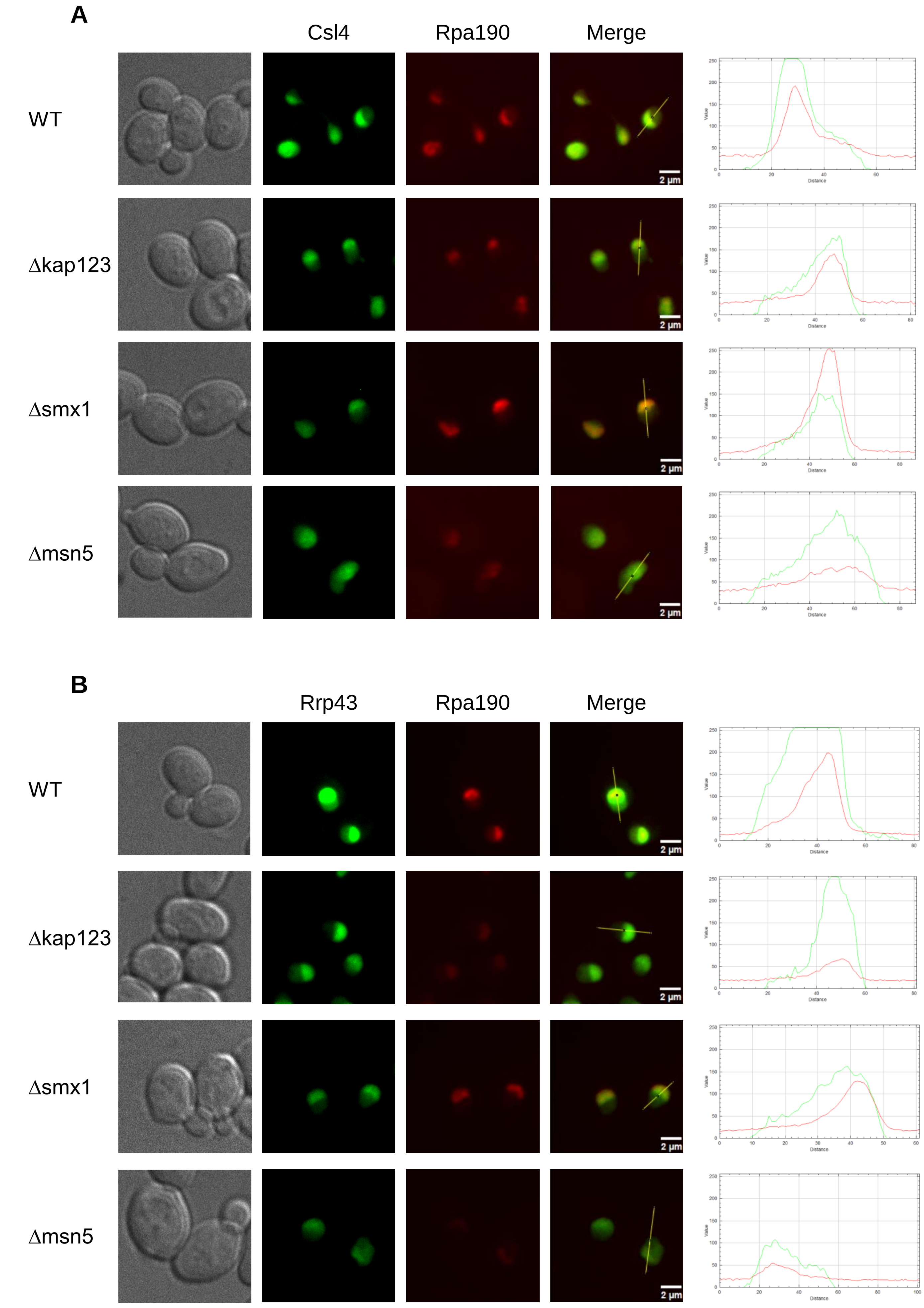

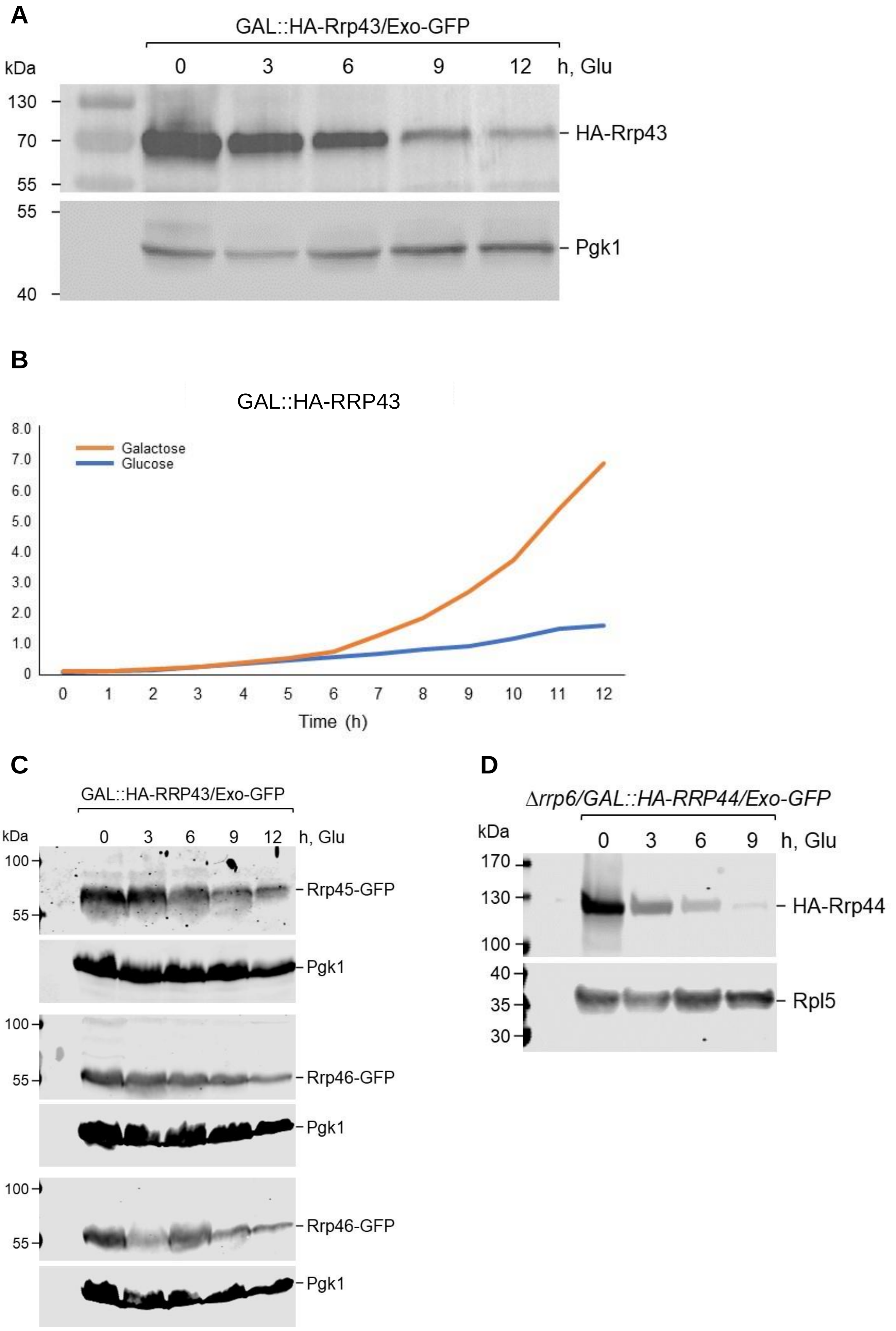

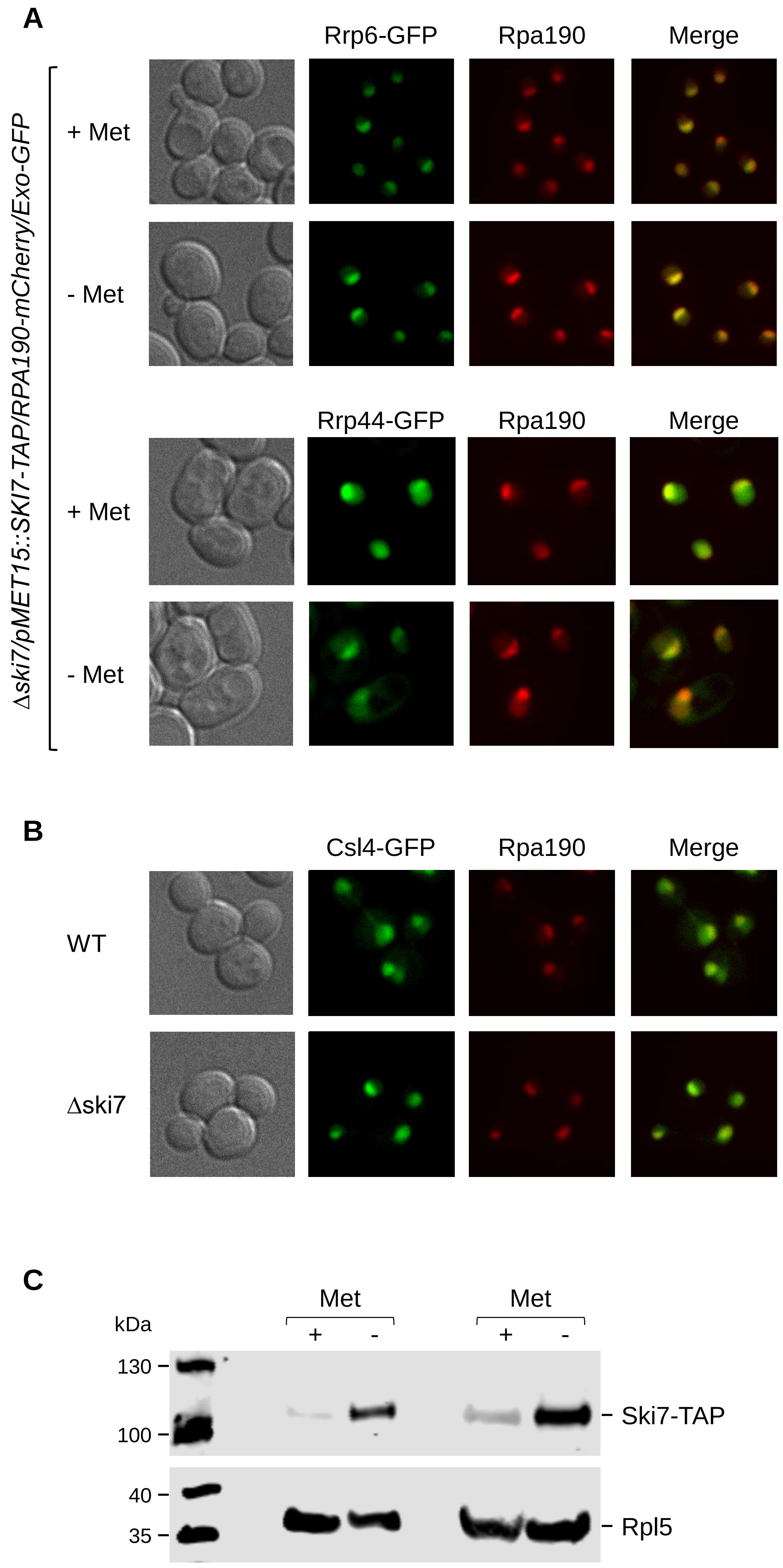

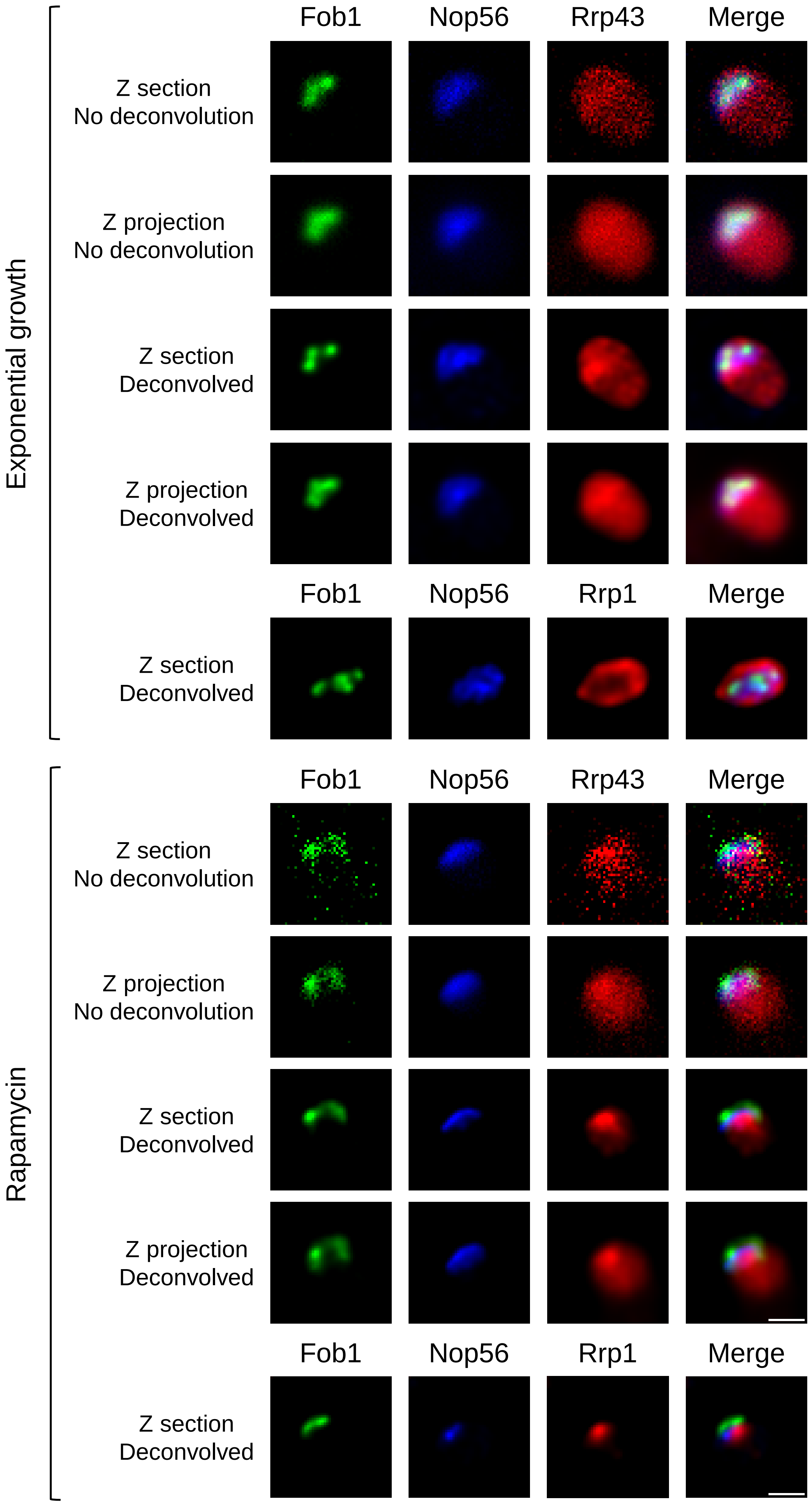

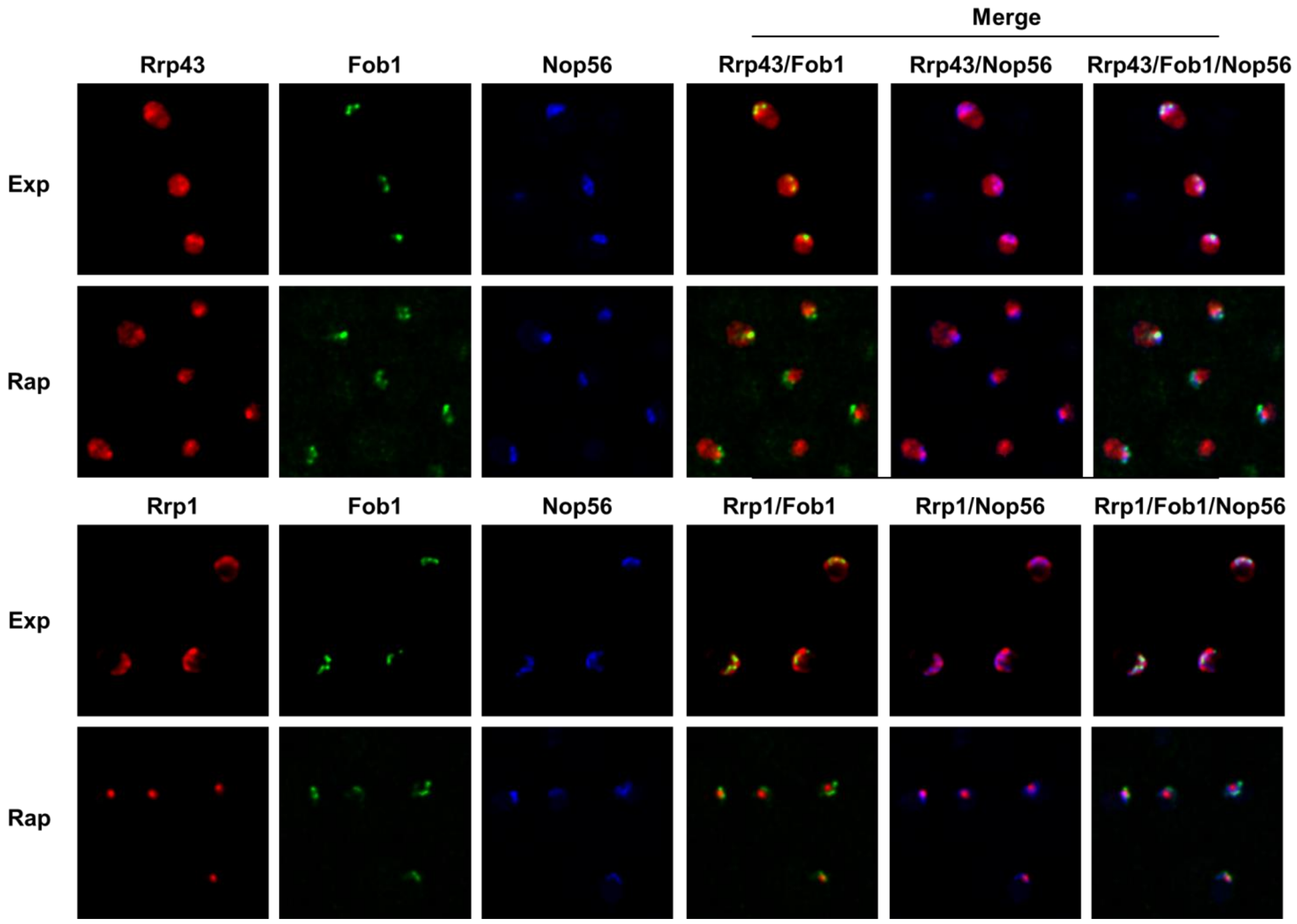
